# Trophoblast paracrine signaling regulates placental hematoendothelial niche

**DOI:** 10.1101/840660

**Authors:** Pratik Home, Ananya Ghosh, Ram Parikshan Kumar, Avishek Ganguly, Bhaswati Bhattacharya, Md. Rashedul Islam, Soma Ray, Sumedha Gunewardena, Soumen Paul

**Author notes:** Co-corresponding authors. Correspondence: Pratik Home; Soumen Paul.

## Abstract

The placenta acts as a major organ for hematopoiesis. It is believed that placental hematopoietic stem and progenitor cells (HSPCs) migrate to the fetal liver to ensure optimum hematopoiesis in the developing embryo. The labyrinth vasculature in a mid-gestation mouse placenta provides a niche for the definitive hematopoietic stem cell (HSC) generation and expansion. It has been proposed that these processes are regulated by a host of paracrine factors secreted by trophoblast giant cells (TGCs) at the maternal-fetal interface. However, the molecular mechanism by which the TGCs regulate the hematoendothelial niche in a developing placenta is yet to be defined. Using a TGC-specific *Gata2* and *Gata3* double knockout mouse model, we show that the loss of GATA2 and GATA3 at the TGC layer leads to fetal growth retardation and embryonic death due to disruptions in the delicate hematopoietic-angiogenic balance in the developing placenta. Using single-cell RNA-Seq analyses, we also show that the loss of GATA factors in the TGCs results in the loss of HSC population within the placental labyrinth and is associated with defective placental angiogenesis. Interestingly, we also found that this TGC-specific GATA factor-loss leads to impaired differentiation and distribution of trophoblast progenitor cells. Our study helps to define the GATA-dependent non-autonomous signaling mechanisms of the primary parietal trophoblast giant cells by which it regulates the delicate hematopoietic-angiogenic balance in the developing placenta.

## Introduction

Trophoblast cells of the placenta establish a vascular connection between the mother and the fetus and express hormones that are essential for the successful progression of pregnancy. Placenta also acts as one of the major organs for hematopoietic stem cell (HSC) generation, and the mid-gestation mouse placenta plays a significant role in the HSC development where it provides a temporary niche for definitive HSC pool. Defective development of placental hematopoiesis and vasculogenesis leads to serious pathological conditions such as preeclampsia and fetal growth retardation (FGR). These disorders result in pregnancy-related complications, maternal, prenatal & neonatal mortality, and affect ∼2–8% of pregnant women worldwide (1, 2). The pathogenesis in preeclamptic patients is believed to be a response of vasculature to abnormal placentation (3). Thus, to define therapeutic modalities against these pregnancy-associated disorders, it is crucial to understand the molecular mechanisms that are associated with the proper development of placental hematopoiesis and vasculogenesis.

Trophoblast cell differentiation in the placenta involves mechanisms by which secreted paracrine factors within the placenta, and from the fetus & the mother regulate embryonic and extraembryonic development (4). Embryonic hematopoietic sites are characterized by the interlinked developments of the vascular and hematopoietic systems. Several studies have shown that the hemangioblasts and hemogenic endothelium act as the presumptive precursors to emerging hematopoietic cells (5). It has been indicated that the definitive hematopoiesis is autonomously initiated in the placenta which subsequently generates HSCs from hemogenic endothelium and provides a niche for expansion of aorta-derived HSCs (6, 7). On the other hand, secreted pro-angiogenic factor, like Placental Growth Factor (Pgf), a member of the Vascular Endothelial Growth Factors (VEGF) family, and anti-angiogenic factors, like Vascular Endothelial Growth Factor receptor-1 (Flt1) and Endoglin (Eng), are expressed in the mouse placenta, indicating autocrine or paracrine actions (8, 9). How these signaling mechanisms influence the development of the maternal-fetal interface are poorly understood.

Several studies have implicated the role of the GATA family of transcription factors in the development of HSCs in other organs, and previously we have shown that GATA2 and GATA3 are involved in the trophoblast development and differentiation (10–13). They are implicated in the regulation of the expression of several trophoblast-specific genes, including prolactin hormone Placental lactogen I (Prl3d1, Pl1) and the angiogenic factor Proliferin (Prl2c2, Plf), which play an essential role in the neovascularization of the placenta (14, 15). Recently we demonstrated that simultaneous knockout of both *Gata2* and *Gata3* in trophoblast lineage severely affects placental development (16). Also, the double gene knock-out resulted in significant developmental defects in the embryo proper leading to very early embryonic lethality (16). These developmental defects were accompanied by severe blood loss in the placenta, yolk sac and the embryo proper. Moreover, we established dual conditional *Gata2* and *Gata3* knockout trophoblast stem cells and used ChIP-Seq and RNA-Seq analyses to define independent and shared global targets of GATA2 and GATA3. We found that several pathways associated with the embryonic hematopoiesis and angiogenesis are targets for both transcription factors.

However, very little is known about how these two master regulators operate in a spatiotemporal manner during placental development and dictate key placental processes like hematopoiesis and angiogenesis.

## Results

### GATA mutation in the trophoblast giant cell layer in mouse embryos display embryonic and extraembryonic hematopoietic defects

Our previous studies have shown that the loss of GATA factors in the trophoblast lineage cells results in the gross phenotypic abnormality in the placenta and the embryo proper in a mouse model (16). The placenta contains distinct layers of differentiated trophoblast cells, each having specialized functions to support a pregnancy. These include Trophoblast Giant Cells (TGC), spongiotrophoblasts (SpT), glycogen trophoblasts, and labyrinthine trophoblasts. Each subclass is characterized by its unique gene expression signatures (17). Thus, it is imperative to analyze how GATA factors act in the different subclasses of trophoblast cells in the context of placental development and function. As TGCs have been reported to express both GATA2 and GATA3, we chose to knockout *Gata2* and *Gata3* together in the TGCs (14). Placental TGCs are marked by the expression of two prolactin family of proteins Prl2c2 (Plf) and Prl3d1 (Pl1). Although Plf expression starts at E6.5 in the parietal TGCs, in the latter part of gestation Plf is also expressed in the spiral artery associated TGCs and canal space associated TGCs (18). Moreover, Plf is expressed in a small population of spongiotrophoblast cells (18). On the other hand, Pl1 expression is restricted to the parietal TGCs only (18). Thus, to restrict GATA deletion exclusively to the parietal TGCs, we used a *Prl3d1^tm1(cre)Gle^* (*Pl1^Cre^*) mouse model, where Cre recombinase is selectively expressed in the Pl1 expressing cells, and established a conditional GATA knockout mouse model *Gata2^f/f^*;*Gata3^f/f^;Pl1^Cre^* (GATA-Pl1 KO) (19, 20). These mice were used to selectively knockout both *Gata2* and *Gata3* in the parietal trophoblast giant cells.

We also developed a murine model in which the trophoblast giant cell layers were fluorescently labeled. We used *Gt(ROSA)26Sor^tm4(ACTB-tdTomato,-EGFP)Luo^/J*, (also known as mT/mG) mouse model, which possesses loxP flanked membrane-targeted tdTomato (mT) cassette and expresses strong red fluorescence in all tissues (21). Upon breeding with Cre recombinase expressing mice, the resulting offsprings have the mT cassette deleted in the cre expressing tissue(s), allowing expression of the membrane-targeted EGFP (mG) cassette located in-frame immediately downstream (**Fig. 1A**).

**Fig. 1:**
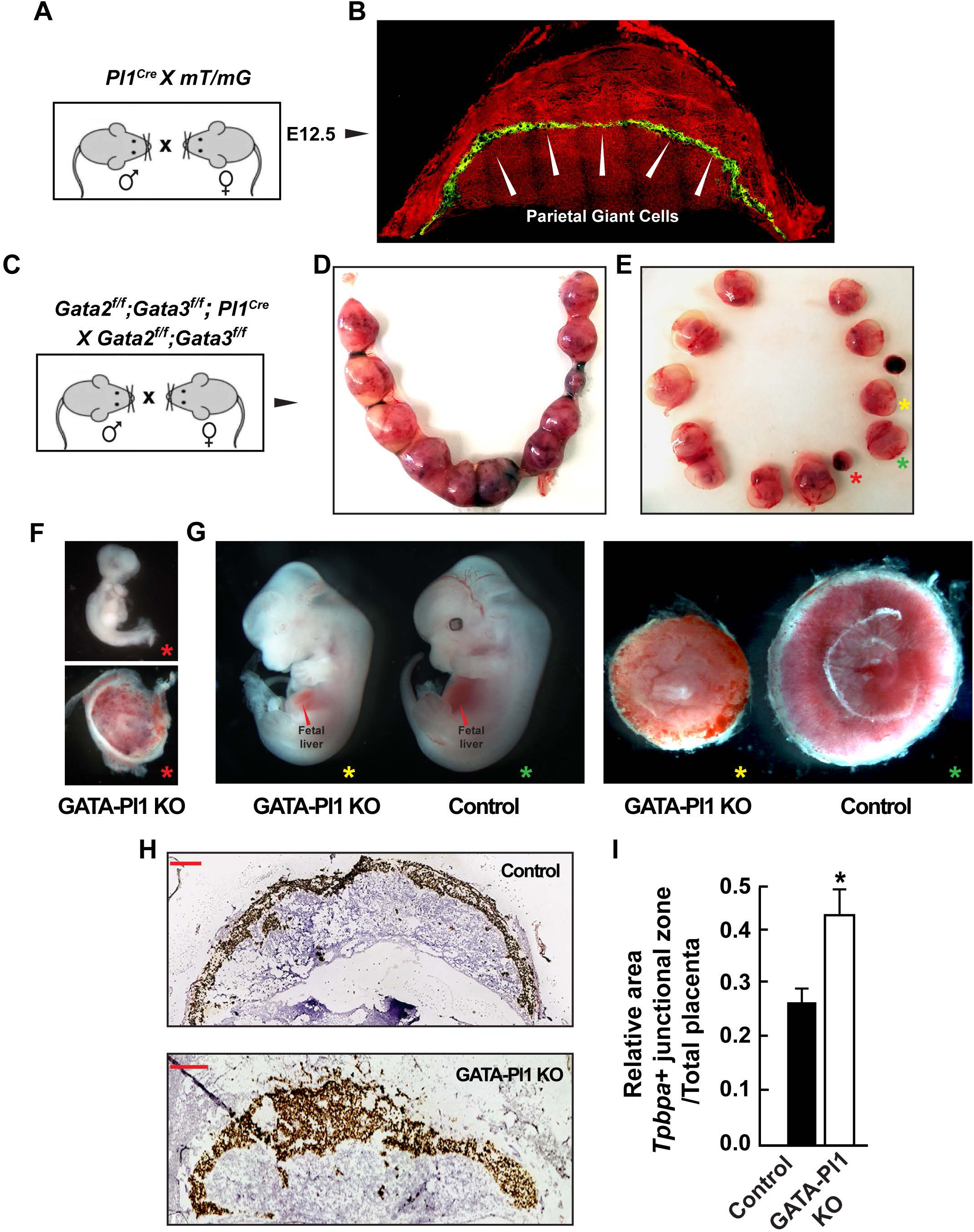
Trophoblast giant cell-specific GATA deletion leads to embryonic lethality and developmental defects. (A) Mating strategy to identify the Prl3d1 positive parietal giant cell layer. (B) Cryosections of the *mT/mG Pl1^Cre^* conceptus show exclusive EGFP expression in the parietal TGCs, indicating the specific nature of the *Pl1^Cre^* expression. (C) Mating strategy to define the importance of GATA factors in the trophoblast giant cells. (D) E12.5 uterine horn harvested from the above mating contains apparent embryonic resorption sites. (E) Isolated conceptuses show severe developmental defects (red asterisk) and size differences compared to the control (green asterisk). (F) Embryos isolated from one of the extremely small conceptuses (red asterisk) reveals gross developmental defects and embryonic death accompanied by small and thin placental tissue. (G) Comparison between a GATA Pl1-KO embryo (red asterisk) and non-Pl1^Cre^ littermate (yellow asterisk) indicates fetal growth reduction, apparent blood loss, and defective vasculature in both the embryo proper and the placenta. Fetal liver of the GATA Pl1-KO embryo also revealed reduced hematopoiesis. (H) RNAScope labeling of implantation sites using *Tpbpa* probe marks the spongiotrophoblast layers. (I) Quantitative comparison of the relative ratio of spongiotrophoblast (SpT) and labyrinth area between the control and GATA Pl1-KO implantation sites(Mean±s.e., n=3, *P≤0.05).

Microscopic analyses of the conceptuses and their cryosections from a cross between *Pl1^Cre^* male and *mT/mG* female confirmed the selective GFP fluorescence in the parietal TGC layers only (**Fig. 1B**). Previous studies have shown that the major expansion of the hematopoietic stem cell (HSC) population in the mouse placenta takes place between E11.5 and E13.5 (22). Thus *Gata2^f/f^*;*Gata3^f/f^;Pl1^Cre^* males were crossed to *Gata2^f/f^*;*Gata3^f/f^* females and the embryos were analyzed between E10.5 and E13.5 (**Fig. C**). Resulting phenotypic abnormalities were observed mostly at E12.5 and E13.5. For all subsequent analyses, E12.5 and E13.5 conceptuses were chosen.

Two major groups of embryos with distinct phenotypic abnormalities were observed in the *Pl1^Cre^* positive embryos (**Fig. 1D, E**). One group showed very early embryonic death and was associated with tiny fetal and placental tissues (**Fig. 1E, F**). The other group showed significant growth retardation, developmental defects, and blood loss in the embryo proper, while their placentae showed significant anomalies in size and thickness and were associated with apparent blood loss (**Fig. 1E, G**). These placentae also showed hemorrhage and apparent loss of vasculature (**Fig. 1G**). These defects were most prominent at E12.5 and E13.5 (**Table 1**). Non-Cre embryos from the same littermates were treated as controls (**Fig. 1E, G**). Along with these gross abnormalities, significant alterations in the placental architecture were observed in the GATA-Pl1 KO samples. A marked increase in the junctional zone (marked by the spongiotrophoblast/ glycogen trophoblast marker *Tpbpa*) (**Fig. 1H, I**) was observed compared to the control.

**Table 1.**

| **Gestation day** | **No. of litters analyzed** | **No. of embryos with Cre** | **No. of embryos without Cre** | **No. of embryos resorbed** | **No. of Cre-positive embryos with growth retardation** |
| --- | --- | --- | --- | --- | --- |
| 10.5 | 6 | 18 | 36 | 2 | 1 |
| 12.5 | 10 | 20 | 56 | 24 | 12 |
| 13.5 | 10 | 18 | 48 | 33 | 10 |

Interestingly, the fetal liver in the GATA-Pl1 KO embryos showed blood loss (**Fig. 1G**). It is speculated that along with the HSPCs from the yolk sac, the placental HSPCs migrate to and seeds fetal liver for hematopoiesis (23). Our results bolster this hypothesis and indicate that the placenta acts as a major contributing partner to the fetal liver hematopoiesis.

A small percentage of *Gata2^f/f^*;*Gata3^f/f^;Pl1^Cre^* embryos escaped the fate and resulted in live-born pups (**Table 1**). These animals showed no apparent phenotypic abnormalities, were healthy and fertile. Genotyping results showed no Cre-mediated excision activities in different tissues (data not shown). For the embryo analyses, we used these males for mating with non-Cre females to restrict the effect of GATA deletion exclusively in the fetal tissue.

Thus, our study showed that the GATA factor-loss in the parietal TGCs of the placenta was sufficient to significantly impair embryonic and extraembryonic growth and also resulted in blood loss and vasculature defects.

### GATA factor functions in parietal giant cells regulate trophoblast progenitor developments

Some of the critical tasks of TGCs are the secretion of autocrine and paracrine factors that are involved in the trophoblast outgrowth and placental development process (24–26). Genetic knockout models targeting TGC specific genes have been shown to affect lineage-specific trophoblast differentiation and thereby results in abnormal development of the placenta (20, 24, 27, 28). To analyze the effect of GATA deletion in the TGCs, we performed Single-cell RNA-Sequencing (scRNA-seq) analyses of E13.5 placenta from a pregnant *Gata2^f/f^*;*Gata3^f/f^* female crossed with *Gata2^f/f^*;*Gata3^f/f^;Pl1^Cre^* male. Two individual GATA-Pl1 KO placentae and two individual control littermate placentae were used for the sequencing. The genotypes were confirmed by PCR using tissue from the embryo proper. The knockout placentae were chosen based on their apparent growth defects and blood loss phenotypes (**Fig. 1G**).

T-distributed Stochastic Neighbor Embedding (t-SNE) plot of the aggregated hierarchical clustering of the four samples revealed 33 distinct clusters (**Fig. 2A**). The clustering of both the control samples showed significant similarity to each other, while significant clustering similarities were observed in both the knock-out samples, indicating a high degree of homology between the placenta of same genotypes (**Sup. Fig. S1A**). A comparison between the control sample and the knockout sample populations showed significant gain in subpopulations defined by individual clusters. While the control samples showed considerable enrichment in clusters 2, 5, 6, 10, 17, 18 and 28, the knockout samples showed enrichment in clusters 1, 3, 7, 9, 14, 20, 21, 23, 29, 30, and 32 (**Fig. 2B**). Trophoblast cell population was identified using keratin 8 (*Krt8*) expression (**Fig. 2C**). Using significant gene expression (p<=0.05) profile of different trophoblast specific genes with a log 2-fold expression of more than zero, we matched major trophoblast gene markers of a mouse placenta with their corresponding clusters (**Sup. Table 1**). While the TGC markers *Prl3d1* (Pl1) and *Prl2c2* (Plf) showed mostly similar expression pattern (clusters 11, 22, 33, 2, 1 and 33, 22, 25, 11, 1 respectively), the spongiotrophoblast marker Trophoblast-specific protein alpha (*Tpbpa*) showed association with clusters 22, 33, 11, 1 and 9 with highest expression in the cluster 22 (**Fig. 2D, Sup. Table 1**). Cathepsin Q (*Ctsq*), a marker for sinusoidal trophoblast giant cells, which line the labyrinth sinusoids and constitute part of the interhemal membrane, showed significant expression associated with cluster 6 (**Fig. 2D, Sup. Table 1**) (31). Epithelial cell adhesion molecule (Epcam), which marks the chorion derived labyrinth progenitors, showed similar expression patterns (clusters 16, 27) in both the control and the KO populations (**Fig. 2D, Sup. Table 1**). Interestingly, prolactin family member *Prl3b1*, which marks both sinusoidal TGCs and parietal TGCs, showed an identical cluster association with *Tpbpa* (**Sup. Fig. S1B, Sup. Table 1**). Both the labyrinth zone markers Syncytin-A (*Syna*) and Syncytin-B (*Synb*) showed significant expression associated with cluster 16 (**Sup. Table 1**). The identity of this cluster was further confirmed by its association with Distal-less homeobox 3 (*Dlx3*) expression (16, 24) (**Sup. Table 1**). Very few cells showed Glial Cells Missing Transcription Factor 1 (*Gcm1*) expression, which confirmed an earlier report that Gcm1 expression reduces significantly after E9.5 (**Sup. Table 1**) (29, 30). We also observed significant expression of Caudal type homeobox 2 (*Cdx2*), a trophoblast stem cell marker, in clusters 20, 26 and 8 (**Sup. Fig. S1B, Sup. Table 1**). Comparative analyses of the control and the KO samples showed significant alterations in these identified trophoblast subpopulations. While *Prl3d1*+ parietal TGCs did not show much change, we observed a marked decrease in the *Prl2c2*+ TGC population in the GATA-Pl1 KO placentae compared to the controls (**Fig. 2E**). Also, *Tpbpa*+ spongiotrophoblast/ glycogen trophoblast population, as well as *Epcam*^Hi^+ labyrinth progenitor population, *Prl3b1*+ secondary giant cells, all showed marked increase in the KO samples, (**Fig. 2E, Sup. Fig. S1B**). Surprisingly, our data also revealed a considerable increase in the *Cdx2*+ cell population in the GATA-Pl1 KO placentae (**Fig. 2E, Sup. Fig. S1B**). However, not all of these cells showed simultaneous expression of other trophoblast stem cell markers E74-like factor 5 (*Elf5*), Eomesodermin (*Eomes*), Estrogen related receptor, beta (*Esrrb*). A fraction of the *Cdx2* expressing cells showed co-expression of *Elf5*, while another fraction was positive for Esrrb. Interestingly, both these fractions were also found to be co-expressing labyrinth marker *Epcam* and *Dlx3*, indicating the role of intermediate progenitor state in the developing labyrinth. We also noticed a significant decrease in the *Ctsq* positive trophoblast cells in the knockout placentae (**Fig. 2D, E**).

**Fig. 2:**
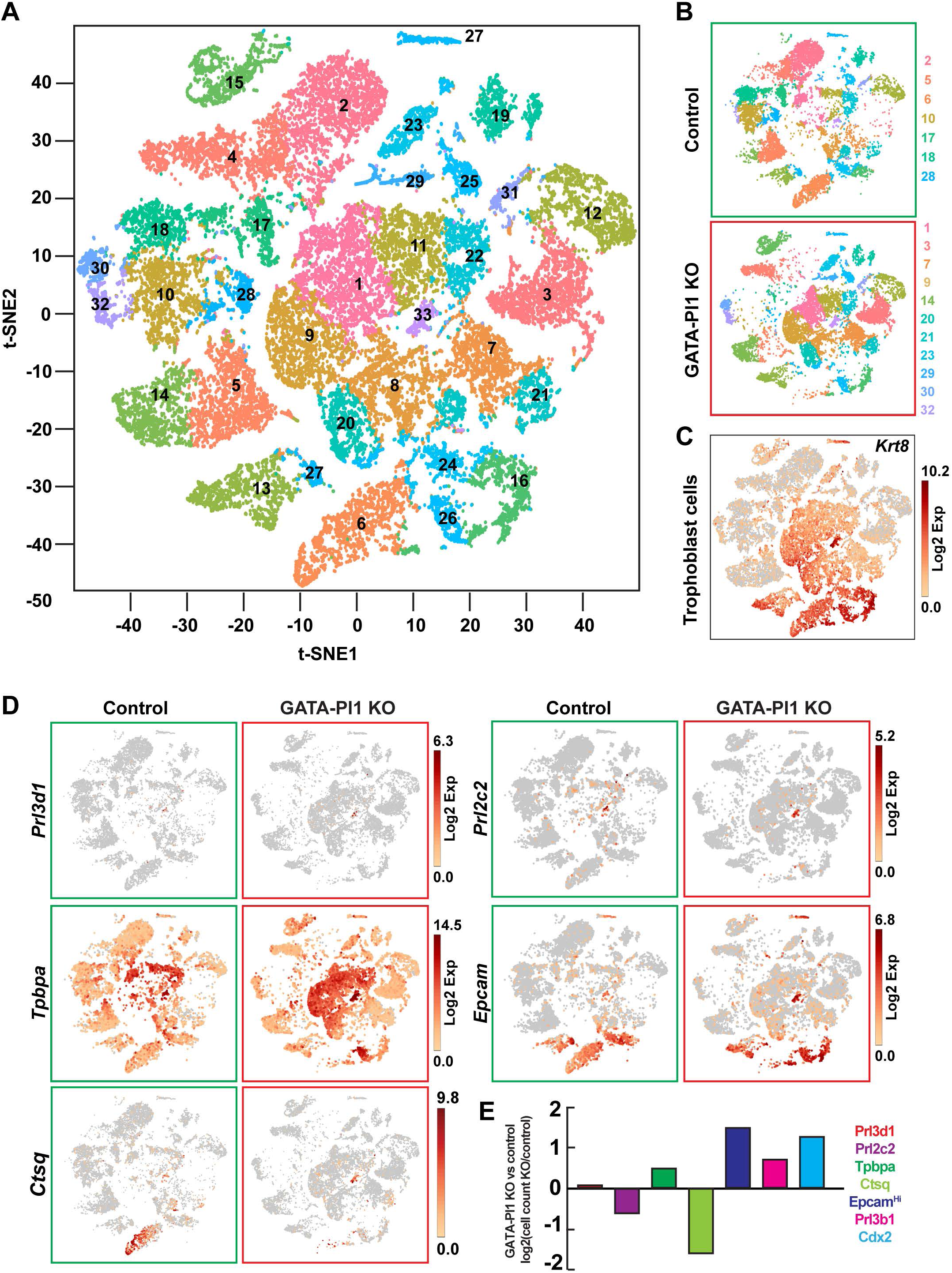
Single Cell RNA-Seq analyses of the TGC-specific GATA factor loss show altered placental trophoblast subpopulation. (A) A t-SNE plot of the aggregate of the hierarchical clustering of 2 control and 2 GATA Pl1-KO placental samples shows 33 distinct clusters. (B) Individual t-SNE plots of the control (aggregate of two control samples) and the KO (aggregate of two KO samples) samples show a gain of trophoblast subpopulations marked by respective clusters on the side. (C) Trophoblast cells are labeled by *Krt8* expressions on a t-SNE plot. (D) Comparative t-SNE plot of cells marked by the expression (log 2-fold expression > 0) of trophoblast lineage markers *Prl3d1*, *Prl2c2*, *Tpbpa*, *Epcam* and *Cdx2* (in the control (green box) vs GATA Pl1-KO (red box) placentae. (E) Quantitative analyses of the relative cell count for the *Prl3d1*, *Prl2c2*, *Tpbpa*, *Cdx2, Epcam*^Hi^, *Prl3b1,* and *Ctsq* positive cells. For all markers except *Epcam*^Hi^, log 2-fold expression > 0; for *Epcam*^Hi^ log 2-fold expression > 3. All cell numbers were normalized using corresponding input cell numbers. Aggregates of two control samples and aggregates of two KO samples were considered for these analyses.

Together, these findings revealed that the parietal TGC-specific loss of GATA factors skews the trophoblast differentiation process and thereby alters the distribution of trophoblast subpopulations in the developing placenta and affects gross placental architecture. Remarkably, these data also indicate a novel method whereby the parietal TGC-specific paracrine signaling dictates the differentiation of the different trophoblast progenitors and regulates the development of distinct placental layers.

### Loss of GATA factors disrupts hematopoietic-endothelial cell lineage segregation and affects fetal hematopoiesis

The placenta is one of the major sites for *de novo* hematopoiesis in an embryo (22). Not only does it support the expansion of the nascent HSC population, but it also protects the HSCs from premature differentiation cues. As the placenta is a known source of a plethora of cytokines and hormones, it is essential to define how these signals promote hematopoietic development in the embryo. Several studies have implicated primary and secondary trophoblast giant cells in secreting paracrine and endocrine factors (31). These include prolactin/ placental lactogen class of hormones (32), interferon (25), vasodilators (33), and anticoagulants (34). Along with them, TGCs have also been shown to secrete angiogenic factors (27, 35, 36). A hallmark of fetal hematopoiesis is the interrelated development of vascular and hematopoietic systems where hemogenic endothelium and hemangioblasts serve as the precursors to the hematopoietic stem cell populations (37).

We used Ingenuity Pathway Analysis (IPA) to compare the physiological functions of the *Prl3d1+* cells between the control and the KO samples using scRNA-seq data. Our results indicated that major physiological functions related to the hematopoiesis and angiogenesis were downregulated in the *Prl3d1+* cells in the GATA-Pl1 KO samples (**Sup. Fig. S2**). These results also bolstered our findings that the GATA factor loss in the parietal TGCs adversely affected embryonic hematopoiesis and angiogenesis.

Placental HSCs are localized in the labyrinth and umbilical blood vessels (38) and are characterized by the surface markers CD34, KIT, and Sca-1 (Ly6A) (39). We used our scRNA-seq data to identify the hematopoietic stem cells in the control compared to the KO samples. We found that the major populations of the HSCs which express both KIT and CD34 are localized mostly in the clusters 15 and 17 (**Fig. 3A, B**). Further analyses of these two clusters revealed that out of these two populations, only the cells in the cluster 15express endothelial cell marker *Kdr*. These suggest that the cells in the cluster 17 represent true hematopoietic stem cell lineage in a developing placenta.

**Fig. 3.**
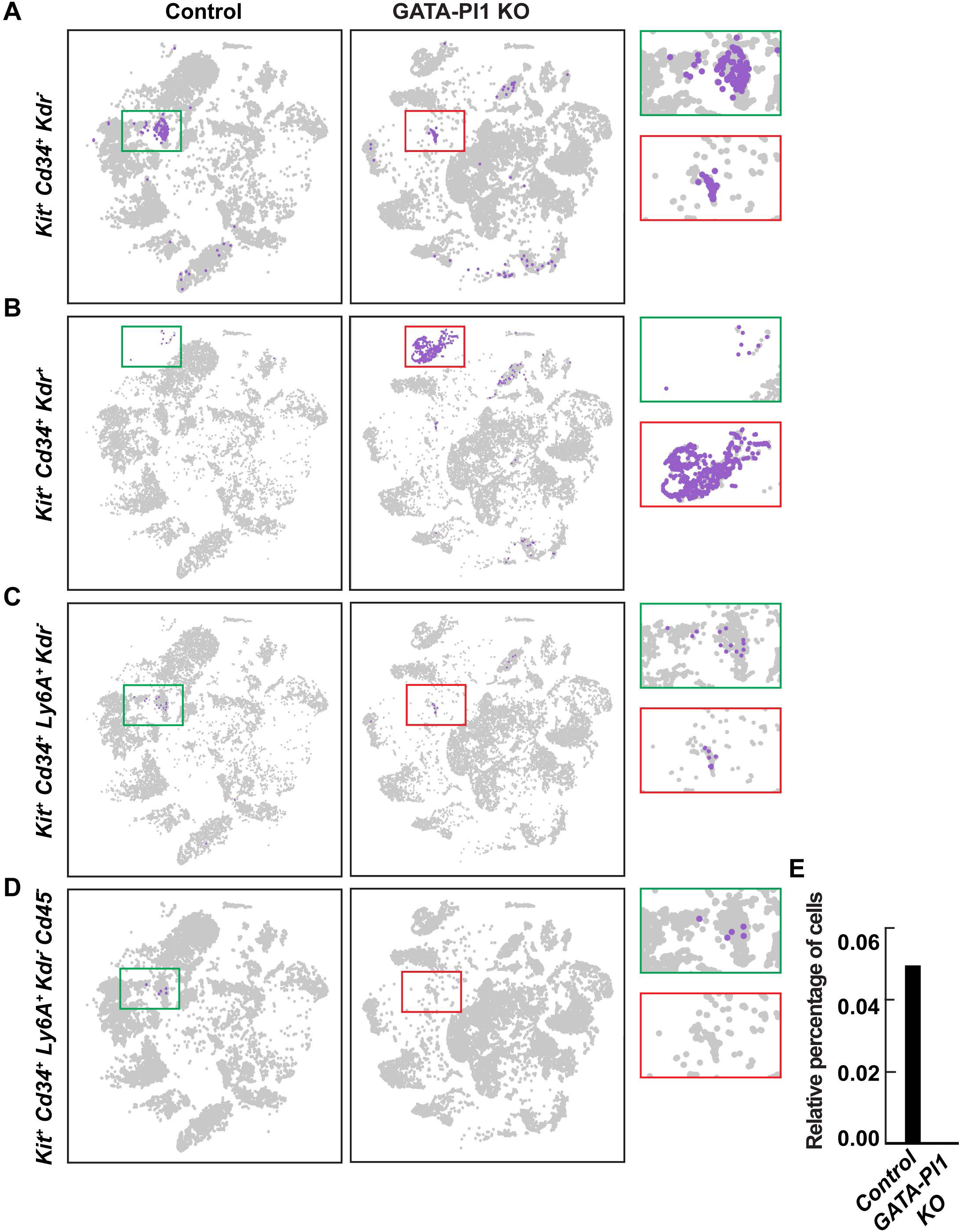
TGC-specific GATA factor function is essential for the differentiation of the hematoendothelial niche. (A) scRNA-seq analyses show cluster 17 to harbor the *Kit*+ *Cd34*+ population of HSCs, which do not express endothelial marker *Kdr*. The t-SNE plot shows a significant reduction in this population in the GATA Pl1-KO placentae compared to the control. (B) Another *Kdr* expressing population of *Kit*+ *Cd34*+ cells (cluster 15) is present in the KO samples, indicating impaired differentiation of the hematoendothelial population upon TGC-specific GATA loss. (C) Another placental HSC marker *Ly6A* (Sca1) is co-expressed by a subset of the *Kit*+ *Cd34*+ *Kdr*-cells present in cluster 17, and this population displays a significant decrease in the KO samples. (D, E) When filtered against the leukocyte marker *Cd45*, these cells in C show a significant decrease in the GATA Pl1-KO placentae compared to the control.

Multiple studies have shown that the embryonic hematopoiesis is intricately connected to the vascular development where hemogenic endothelium, a part of the vascular endothelial cells gives rise to the definitive hematopoietic precursors during mammalian development (40–44). Thus, the genetic signature of the cells in cluster 17, which is unique to the knockout samples, indicates a hematoendothelial cell population that still retains the HSC lineage markers (**Fig. 3B**). This cell population was also found to express hematoendothelial markers *Cdh5*, *Icam2*, *Cd40*, confirming the bipotent nature of these cells (45–47). As a high expression of the arterial-specific marker Delta-like, 4 (*Dll4*) is essential for the segregation of the endothelial lineage from the hematopoietic lineage (48, 49), we looked at the *Dll4* expression in this subset, and found that this cluster expresses a high level of *Dll4*. Thus, the loss of GATA factors in the parietal TGCs skews the hematopoietic-endothelial lineage specification in a developing placenta and leads to the arrest of hematoendothelial progenitor population compared to the control.

We further analyzed these cells for the expression of *Sca-1*, another HSC marker, and *Cd45*, a differentiated hematopoietic marker, and found that while the small HSC population in cluster 17 express *Sca-1*, they do not express *Cd45* (**Fig. 3C, D**), confirming their undifferentiated HSC identity. Further analysis Quantitative analysis showed a severe reduction in the HSC population in the GATA-Pl1 KO placentae compared to the control (**Fig. 3E**).

As the vascular labyrinth of a developing placenta harbors HSC population, we also tested the loss of the HSC population in the placental labyrinth by immunohistochemistry. E12.5 and E13.5 placental sections were stained with an anti-KIT antibody, while the placental layers were co-stained using an anti-pan-cytokeratin antibody. The control placenta showed several clusters of KIT^+^ cells in the labyrinth, while consistent with our scRNA-seq data, the KO placenta showed severe loss of KIT^+^ cell population (**Fig. 4A**).

**Fig. 4:**
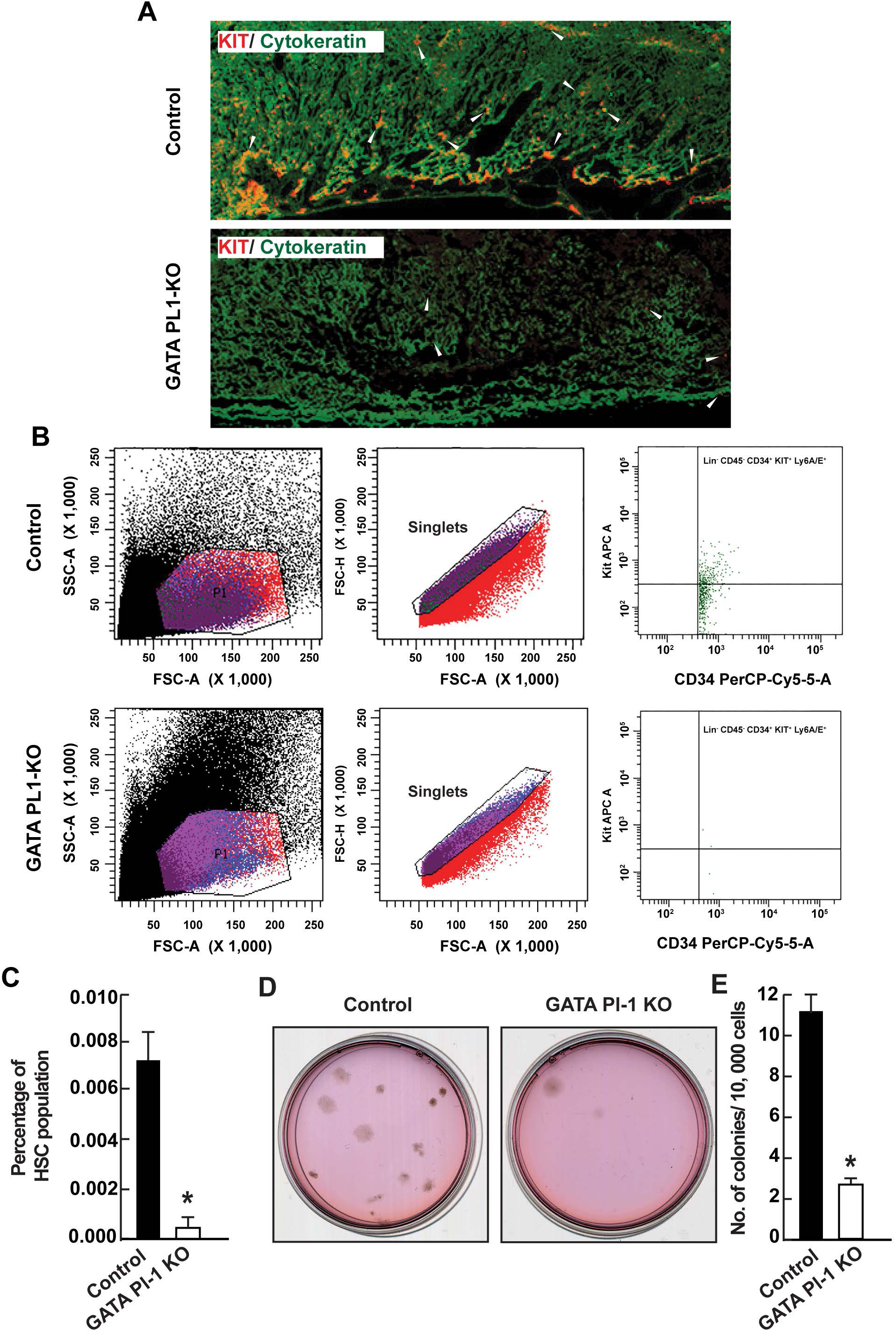
GATA factor loss in the TGCs results in the loss of the hematopoietic progenitor population. (A) KIT staining of the implantation sites shows the loss of KIT-positive HSC clusters in the junctional zone and the labyrinth of the KO placenta compared to the control. The anti-pan-cytokeratin antibody was used to mark the trophoblast layers. White arrows indicate KIT-positive cell clusters around the vasculature. (B) Flow analysis data shows a significant reduction in the HSC population in the knockout placenta. E12.5 placental samples from GATA Pl1-KO and control littermate placentae were subjected to flow analysis. CD34^+^ KIT^+^ Ly6A/E^+^ cells were counted against Lin^-^ CD45^-^ cells. (C) Quantitation of the Lin^-^ CD45^-^ CD34^+^ KIT^+^ Ly6A/E^+^ population reveals significant loss of the HSC population in the KO sample (Mean±s.e., n=3, *P≤0.05). (D) Micrographs show the formation of hematopoietic cell colonies on a methylcellulose plate. Colonies included were of CFU-G/M, BFU-E, and CFU-GEMM nature. (E) Quantitation of the number of colonies in the control vs. GATA Pl1-KO samples (Mean±s.e., n=3, *P≤0.05).

We also used three surface markers CD34, KIT and Sca-1 (Ly6A), which together define the placental HSCs, as the determinant for the long-term reconstituting (LTR) HSC population and screened them against the population expressing CD45, which is a pan-hematopoietic surface marker and appear on the more mature HSCs (50), and a cocktail of lineage markers (Lin). We used placental single-cell suspension from control and GATA-Pl1 KO placentae and subjected them to flow analysis where KIT^+^ CD34^+^ Ly6A/E^+^ cells were selected against Lin^-^ CD45^-^ expressing cells. In the GATA-Pl1 KO placental samples, the Lin^-^ CD45^-^ KIT^+^ CD34^+^ Ly6A/E^+^ cell population was significantly reduced compared to the control (**Fig. 4B**). Compared to the total number of cells, the percentage of the HSC population in the KO placenta (0.001±0.000) was significantly lower than that in the control population (0.006±0.001) (**Fig. 4C**).

Next, we evaluated the differentiation potential of the GATA-Pl1 KO placental HSCs by using colony-forming assay. Placental single-cell suspension from the control and GATA-Pl1 KO placentae were plated on methylcellulose medium containing a cocktail of Stem Cell Factor (SCF), IL-3, IL-6, and Erythropoietin. The placental samples gave rise to mostly granulocyte and mixed lineage erythroid, macrophage colonies (**Fig. Sup. S3**). We observed a significant reduction in the number of colonies generated from the GATA-Pl1 KO placentae compared to the control (**Fig. 4D, E**).

Collectively these data prove that the GATA factor KO in the trophoblast giant cells negatively affects hematopoiesis in the placenta and results in the apparent blood loss phenotype. The loss of the HSC population was also accompanied by the incomplete segregation of the hematopoietic and endothelial lineage, indicating a loss of signaling network that balances and fine-tunes the hematopoietic versus endothelial cell lineage development in the placenta.

### TGC-specific loss of GATA factors affects embryonic vasculature development

Embryonic hematopoiesis and angiogenesis are tightly linked. Hemogenic endothelium in the vascular labyrinth gives rise to both the endothelial cell population as well as the hematopoietic cell population. As our study revealed a defective hematopoietic and endothelial lineage segregation due to TGC-specific GATA factor loss, we used our GATA-Pl1 KO placentae model to test placental angiogenesis. Anti-CD31 (PECAM-1) antibody, which marks early and mature endothelial cells, was used to stain the vasculature in the placental sections at E13.5. Compared to the control samples, the KO placentae showed gross disruption of the blood vessel architecture in the labyrinth. The KO placental samples displayed defective branching, unlike the complex placental vasculature branching observed in the control samples (**Fig. 5A, B**).

**Fig. 5:**
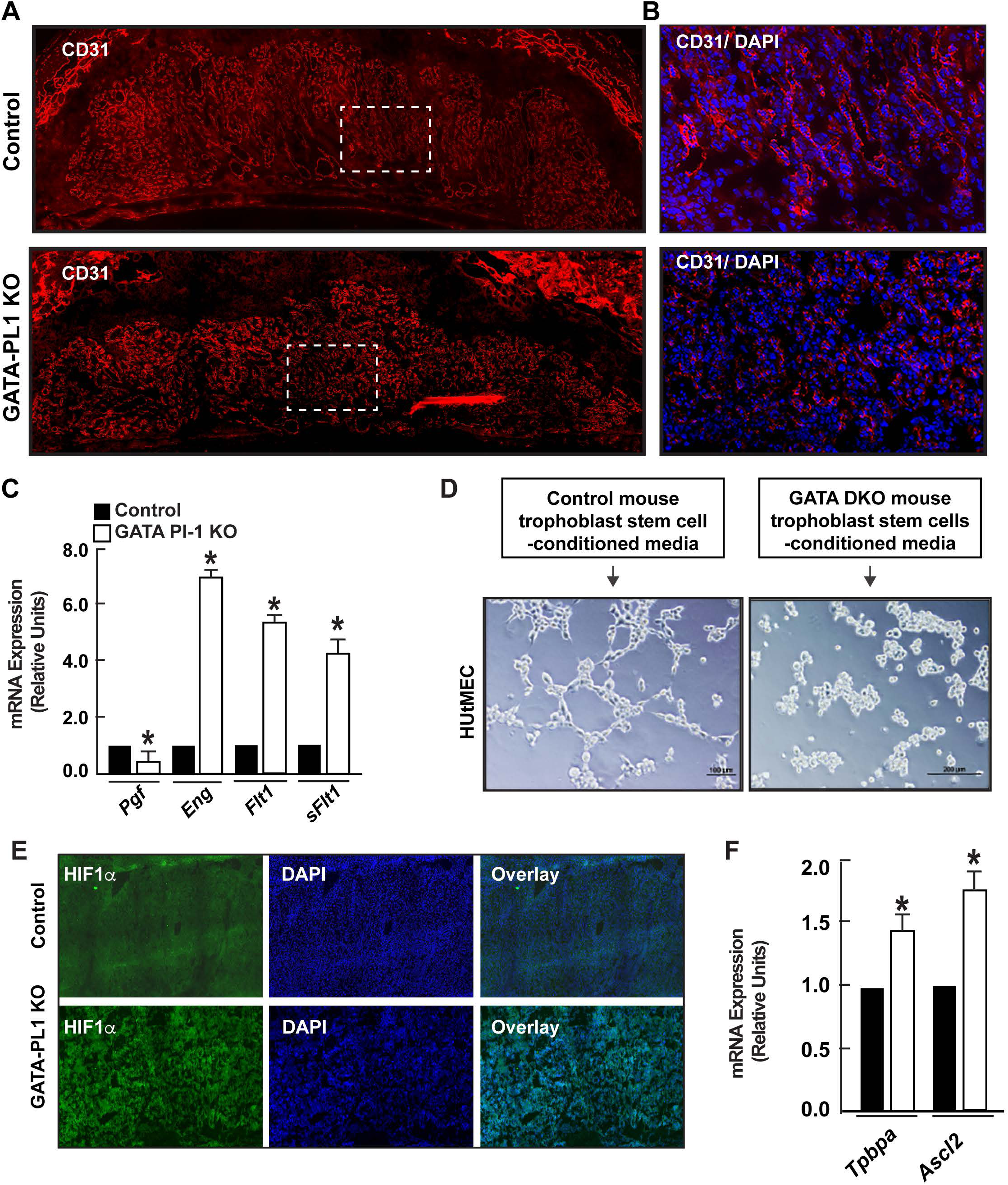
GATA factor-functions in the trophoblast giant cells are essential for the vasculature development at the maternal-fetal interface. (A) Immunostaining of the control and GATA Pl1-KO placenta with the endothelial cell marker CD31 shows abnormal vasculature development upon GATA factor loss in the TGC layer. (B) GATA Pl1-KO placental labyrinth is associated with the defective branching pattern of the blood vessels. (C) mRNA expression analyses show downregulation of pro-angiogenic factor *Pgf* expression compared to the upregulation of anti-angiogenic factors *Eng*, *Flt1*, and *sFlt1* in the KO placenta compared to the control indicating a perturbation of the delicate balance needed between the angiogenic and anti-angiogenic factors for proper vascular development (Mean±s.e., n=3, *P≤0.05). (D) In the presence of conditioned medium from GATA2/GATA3 double knockout (GATA DKO) trophoblast stem cells, Human Uterine Microvascular Endothelial Cells (HUtMEC) fail to form vascular ring-like structures in a matrigel based vascular tube formation assays unlike when treated with conditioned medium from control trophoblast stem cells. (E) Immunostaining data shows a high level of HIF1α in the GATA Pl1-KO placenta compared to the control. (F) mRNA expression analyses show upregulation of *HIF1α* is associated with significantly increased expression of trophoblast differentiation markers *Tpbpa* and *Ascl2* in the KO placenta (Mean±s.e., n=3, *P≤0.05).

Embryonic angiogenesis relies on a delicate balance of several key pro-angiogenic and anti-angiogenic factors (51–54). Among them, pro-angiogenic placental growth factor (*Pgf*), anti-angiogenic factors soluble TGF-β1 receptor Endoglin (*Eng*), and soluble Vascular Endothelial Growth Factor (VEGF) receptor Flt1 (*Fflt1*) have been shown to play essential roles in the placental vascularization and angiogenesis (55–58). We tested our knockout samples for the mRNA expression of these genes. Our results revealed significant downregulation of proangiogenic *Pgf* while significant upregulation of antiangiogenic *Eng*, *Flt1* and its soluble isoform *sFlt1* was observed in the GATA-Pl1 KO placentae compared to the control (**Fig. 5C**).

To functionally test this increase in the *sFlt1* level, we performed matrigel based vascular tube formation assays using Human Uterine Microvascular Endothelial Cells (HUtMEC). Harvested conditioned media from the *Gata2/Gata3* double knockout (GATA DKO) trophoblast stem cells and control trophoblast stem cells described in our earlier study (16) were added to the assays individually. Interestingly, HUtMECs in the presence of the conditioned media from the control trophoblast stem cells, readily formed tubular structure while they failed to do so in the presence of conditioned media from GATA DKO trophoblast stem cells (**Fig. 5D**).

These findings indicate that the loss of GATA factors in the TGC layer results in the disruption of the delicate balance between the secreted angiogenic and antiangiogenic factors, which in turn prevent proper development of the labyrinth vasculature in the placenta.

Multiple studies have implicated HIF family of proteins, HIF1α, and HIF2α, in the trophoblast differentiation process, and they have been shown to promote spongiotrophoblast differentiation in the placental development (59, 60). HIF family of proteins are also intricately connected to angiogenesis. On the one hand, they have been shown to be the master regulators of angiogenesis, while on the other hand, defective vasculature leads to hypoxia which in turn stabilizes HIF1α (61, 62). Since TGC-specific GATA factor-loss resulted in a vasculature defect, we designed experiments to study the role of HIF1α in the knockout placenta. HIF1α immunostaining of the cryosectioned implantation sites showed significantly stronger HIF1α expression in the GATA Pl1-KO placenta compared to the control. Thus, the defective vasculature in the knockout embryos were associated with the upregulation and stabilization of HIF1α (**Fig. 5E**).

It has been demonstrated that the HIF1α overexpression in mice leads to fetal IUGR and abnormalities in the developing placenta through the upregulation of *sFlt1* (63). As we have discussed above, GATA Pl1-KO samples showed elevated expression of *sFlt1*, indicating a possibility that the fetal growth retardations observed in the GATA Pl1-KO embryos are regulated by this HIF1*α−*sFlt1 regulatory axis.

HIF1α stabilization also induces spongiotrophoblast markers *Tpbpa* and *Ascl2* (64) and promotes spongiotrophoblast differentiation (65). mRNA expression analyses of the knockout placenta revealed a significant increase in both *Tpbpa* and *Ascl2* level (**Fig. 5F**), leaving scope for further evaluation of the HIF1α feedback loop in GATA Pl1-KO placental development.

### GATA loss alters trophoblast giant cell-mediated paracrine signaling

TGCs are known to be a major source of autocrine and paracrine factors in the placenta (18, 66). These factors, in turn, influence the development of the placenta as well as regulate numerous placental functions. We used our *Gata2^f/f^*;*Gata3^f/f^;mT/mG* mouse model to analyze the paracrine signaling in the light of GATA loss in the TGCs. As described above, E12.5 conceptuses from a cross between *Gata2^f/f^*;*Gata3^f/f^;Pl1^Cre^* and *Gata2^f/f^*;*Gata3^f/f^*;*mT/mG* were harvested and chosen for strong GFP fluorescence indicating the presence of Pl1^Cre^ (**Fig. 6A**). The GFP-positive cells were sorted using FACS (**Fig. 6B**). *Pl1^Cre^*;*mT/mG* conceptuses were used as control. mRNA samples were prepared from these cells and were used for subsequent qPCR analyses. As TGCs are marked by the expression of *Prl3d1* and *Prl2c2* and it has been shown that these two factors are regulated by GATA factors (16, 67), we tested their expression in the GATA Pl1-KO samples. Both *Prl3d1* and *Prl2c2* mRNA prepared from these cells showed a significant reduction in their expression levels compared to the control (**Fig. 6C**). Furthermore, we tested several endocrine and paracrine factors secreted by the TGCs that are important for hematopoiesis and angiogenesis.

**Fig. 6:**
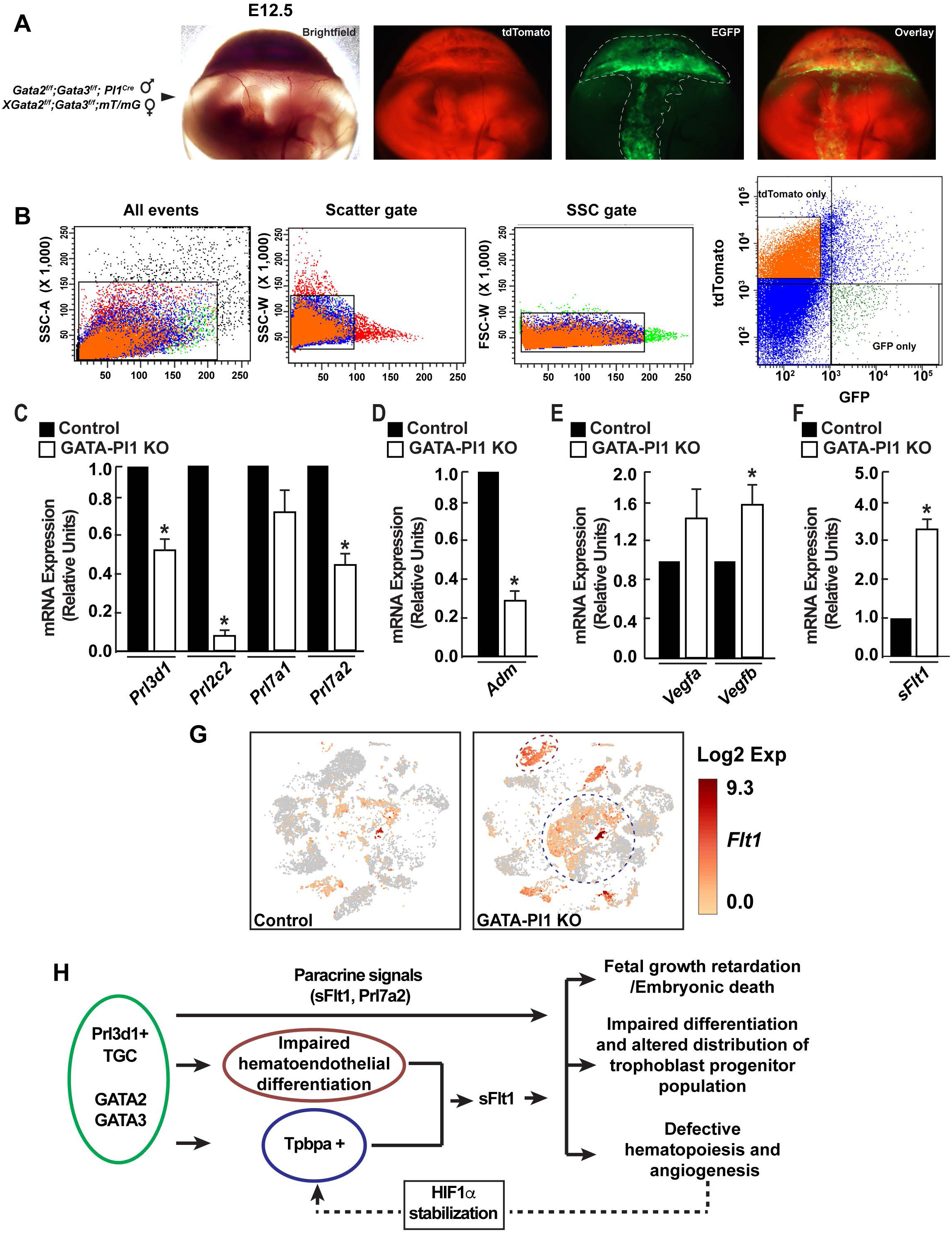
The GATA factors regulate parietal TGC-specific paracrine factors and help maintain hematopoietic-angiogenic balance. (A) Isolated E12.5 conceptus from a *Gata2^f/f^*;*Gata3^f/f^;Pl1^Cre^* (male) X *Gata2^f/f^*;*Gata3^f/f^*;*mT/mG* (female) show distinct EGFP expression in the parietal TGC layer. (B) EGFP+ GATA Pl1-KO TGCs were FACS purified from the *Pl1^Cre^* placentae. EGFP+ TGCs isolated from *Pl1^Cre^; mT/mG* placentae were used as controls. (D) Significant downregulation of *Prl3d1*, *Prl2c2*, and *Prl7a2*, while no significant change in the *Prl7a1* expression is associated with the GATA Pl1-KO TGCs (Mean±s.e., n=3, *P≤0.05). (D) Vasodilator hormone Adrenomedullin (*Adm*) expression shows a significant reduction in the KO TGC samples compared to the control (Mean±s.e., n=3, *P≤0.05). (E, F) mRNA expression reveals significant upregulation of *Vegfb* and antiangiogenic *sFlt1* in the KO TGCs, while no significant change is associated with *Vegfa* (Mean±s.e., n=3, *P≤0.05). (G) t-SNA plot shows a marked increase in *sFlt1* expressing cells in the GATA Pl1-KO placentae, where the major source of the *sFlt1* is the *Tpbpa*+ spongiotrophoblast layer (circled by blue dashes) and the hematoendothelial population in cluster 17 (circled by purple dashes). (H) Our proposed model shows that the GATA factors in the *Prl3d1*+ TGCs function regulate proper embryonic growth, placental cell differentiation, distribution, and maintain delicate hematopoietic-angiogenic balance. These regulations are achieved by the paracrine signals sFlt1 and Prl7a2 and also in parallel by the stabilization of HIF1α. Stabilized HIF1α alters the *Tpbpa*+ spongiotrophoblast layer, which in turn contributes to the increased antiangiogenic *sFlt1* level in the developing placenta.

Among the prolactin like protein family members secreted by the TGCs, Prl7a1, and Prl7a2 are known to regulate erythroid proliferation and differentiation as well as megakaryocyte differentiation (68, 69). Our qPCR analyses revealed significant reductions in *Prl7a2,* but not in *Prl7a1* expression in the TGCs from GATA Pl1-KO placentae (**Fig. 6C**). These results indicate to the possibility that the loss of the HSC population in the double knockout placenta was partially due to the downregulation of *Prl7a2* in the TGCs.

TGCs also secrete the hormone Adrenomedullin (*Adm*), a vasodilator implicated in the placental angiogenesis (70, 71). A low level of adrenomedullin contributes to the preeclamptic human placenta (72). We tested the level of *Adm* in the double knockout TGCs and found that the Adm level was significantly downregulated (**Fig. 6D**).

Additionally, our analyses of the VEGF family members in the GATA Pl1-KO TGCs revealed significant upregulation of *Vegfb* while *Vegfa* showed no significant difference compared to the control (**Fig. 6E**). We also observed significant upregulation of s*Flt1* in the knockout samples (**Fig. 6F**). Interestingly, our scRNA-seq data revealed that along with the arrested hematoendothelial progenitor population, *Tpbpa*+ spongiotrophoblast cells were a significant source of sFlt1 in the knockout placenta (**Fig. 6G**), which is consistent with an earlier report (8). Thus, our findings reveal that the loss of GATA factors influenced the level of Flt1 in the knockout placentae in two distinct ways. On the one hand, it increases the sFlt1 level in the GATA Pl1-KO TGCs while, on the other hand, it increases the number of *Tpbpa+* spongiotrophoblast cells and in turn increases the sFlt1 level in the knockout placentae.

Overall, these findings implicate TGC-specific GATA2 and GATA3 to be not only critical regulators of placental development but also essential players for maintaining the delicate balance between the hematopoietic and endothelial lineage segregation in the developing placenta. Thus, we propose a model where GATA2 and GATA3 in the *Prl3d1*+ parietal TGCs control TGC-specific paracrine signaling and thereby directly regulates the differentiation of the hematoendothelial niche in the placental labyrinth and promotes the differentiation of the spongiotrophoblast layer (**Fig. 6H**). Upregulated sFlt1 from the GATA-null parietal giant cells, enhanced spongiotrophopblast layer and the impaired hematoendothelial niche, results in the fetal growth retardation, altered trophoblast progenitor differentiation program and defective hematopoiesis and angiogenesis. This impaired placental angiogenesis contributes to the upregulation and stabilization of HIF1α, which in turn affects the *Tpbpa*+ spongiotrophpblast differentiation through the alteration of sFlt1 level in a negative feedback loop manner (**Fig. 6H**).

## Discussion

GATA factor function in trophoblast differentiation and function has been subjected to numerous studies. Previously, we showed how GATA2 and GATA3 have overlapping functions during placental development, where the simultaneous loss of both the factors in all trophoblast cells leads to early embryonic death and placental defect with impaired hematopoiesis (13, 16). It also revealed the loss of placenta layers, including the labyrinth region. Interestingly, the pan-trophoblast-specific knock out did not show any change in the trophoblast giant cell layer at the junctional zone. So, in this study, we chose to delete GATA2 and GATA3 in the *Prl3d1* expressing parietal giant cell layer of the placenta.

As the placenta is a complex tissue consisting of diverse trophoblast subtypes as well as hematopoietic and endothelial cell populations, it is of utmost importance to categorize these cellular subtypes and define cell-cell interactions at the single-cell resolution. Our Single Cell RNA-Seq data not only revealed the genetic signatures of the major trophoblast subtypes but also showed how the GATA-factor loss altered the differentiation of the progenitor population and thereby altered the distribution of cells at different placental layers. Data from our *Gata2^f/f^*;*Gata3^f/f^;Pl1^Cre^* mouse models showed that the loss of the GATA factors in the parietal TGC population severely reduced the placental size and also led to fetal growth retardation with significant phenotypic abnormality between E12.5 and E13.5. Importantly, we observed significant ultrastructural changes in the GATA-Pl1 KO placentae compared to the control. Molecular and morphological analyses revealed gross redistribution of different trophoblast progenitor cells including *Prl2c2* expressing giant cells, *Tpbpa* expressing spongiotrophoblast progenitor, *Prl3b1* expressing secondary giant cell progenitors. While the number of *Epcam* expressing labyrinth progenitor cells showed a marked increase, we did not observe any significant enlargement of the labyrinth regions in the GATA-Pl1 KO placentae when normalized against the total placental area. Although we noticed an increase in the Cdx2 positive cell population, these cells did not express other trophoblast stem cell markers *Elf5*, *Esrrb* and *Eomes*. As the number of *Cdx2* positive cell population decreases significantly with the maturation of the placentae, it would be interesting to find out the role of these *Cdx2* expressing cells in the mid-gestation mouse placenta. Taken together, these findings reveal how different layers of trophoblast cells interact during placental development and influence the differentiation of trophoblast progenitors.

TGCs in the placenta secrete a lot of cytokines and hormones, which are not only crucial for trophoblast differentiation and placental development but are also critical for placental angiogenesis and hematopoiesis. Our study also characterized how the TGC-specific GATA factors dictate TGC-specific paracrine signaling and in turn, regulate placental hematoendothelial niche. Curiously, the loss of TGC-specific GATA factors resulted in a unique cell population marked by the expression of *Kdr*, as well as *Kit*, *Cd34,* and *Ly6A*. The expression of both the endothelial gene and hematopoietic stem cell markers revealed the emergence of a hematoendothelial cell cluster. The complete absence of this cell population in the control placentae also indicated arrested differentiation of these cells in the absence of TGC-specific GATA factors. Our scRNA-seq data also revealed the true hematopoietic stem cell population in the developing mouse placenta.

The GATA-Pl1 KO placentae were also characterized by the lack of proper angiogenic branching in the labyrinth. Our observation about the increased level of anti-angiogenic factors *Eng* and *sFlt1* and a decrease in the pro-angiogenic *Pgf* level gives us a possible explanation about the angiogenic defect. Further analyses showed the increased *Tpbpa* expressing cells and the immature hematoendothelial cells to be the main contributors to the increased level of *Flt1*. As the *Ctsq* expressing sinusoidal TGCs regulate the maternal sinusoid development, the concomitant decrease in the Ctsq expressing cells in the GATA-Pl1 KO could also contribute to the impairment of the labyrinth vascular development. We speculate that this loss in the surface area of exchange may result in the reduction of the nutrient supply and finally result in fetal growth retardation.

Mouse embryogenesis involves vascularization on both the placental side as well as the decidual side. TGCs secrete a lot of angiogenic and anti-angiogenic paracrine factors, and in turn, regulate the decidual vascularization. However, how these factors dictate decidual vascularization, is poorly understood. It would be interesting to study how TGC-specific loss of GATA factors affect decidualization and associated vascular development. In this aspect, our TGC-specific GATA knockout mouse model holds great promise to help investigate the paracrine signaling from the TGC layers involved in the decidual angiogenesis.

GATA2 and GATA3 are evolutionarily conserved among mammals, and they are expressed in the human placenta. However, unlike the mouse placenta, human placental layers do not have trophoblast giant cells. In the absence of defined TGC subtypes, it is impossible to analyze the role of GATA factors in the placental development in humans. Moreover, ethical and logistical issues make it impossible to examine the role of GATA factors in human placental development *in vivo*. Nonetheless, the recent establishment of a human trophoblast cell line has presented us with a new tool to examine the role of GATA factors in human placental development. As these cells can be readily differentiated *in vitro* to extravillous trophoblast and syncytiotrophoblast subtypes, they open up the possibility of serving as models to study the role of GATA factors in the context of human placental development.

## Materials and methods

### Cell culture and reagents

Mouse trophoblast stem cells (TSCs) were cultured with FGF4, Heparin, and MEF-conditioned medium (CM) according to the protocol (73). *Gata2* and *Gata3* floxed alleles were efficiently excised from *Gata2^f/f^;Gata3^f/f^;UBC-cre/ERT2* (GATA DKO) TSCs by culturing the cells in the presence of tamoxifen (1 μg/ml) (16). Conditioned media was harvested upon removal of tamoxifen and was used for subsequent experiments.

### Generation of conditional knockout mice strains

All procedures were performed after obtaining IACUC approvals at the Univ. of Kansas Medical Center. Female *Gata2flox/flox* (*Gata2^f/f^*) mice (74) were mated with *Prl3d1tm1(cre)Gle* (*Pl1^Cre^*) male in order to generate *Gata2^f/+^;Pl1^Cre^*. In the next step, *Gata2^f/+^;Pl1^Cre^* female mice were bred with *Gata2^f/+^;Pl1^Cre^* males to generate *Gata2^f/f^;Pl1^Cre^*. Similarly female *Gata3flox/flox* (*Gata3^f/f^*) mice (75) were used to generate *Gata3^f/f^;Pl1^Cre^*. In the next step, *Gata2^f/f^;Pl1^Cre^* and Gata3^f/f^;*Pl1^Cre^* mice were crossed to generate Gata2^f/+^;*Gata3^f/+^*;*Pl1^Cre^*. Later *Gata2^f/+^;Gata3^f/+^;Pl1^Cre^* males and females were crossed to generate *Gata2^f/f^;Gata3^f/f^;Pl1^Cre^* strain. Further crosses with Gt(ROSA)26Sortm4(ACTB-tdTomato,-EGFP)Luo/J, (also known as mT/mG) mouse strain was used to establish *Gata2^f/f^;Gata3^f/f^;mT/mG* mouse line.

### Embryo harvest and tissue isolation

Injected animals were euthanized on at desired day points, as indicated in the main text. Uterine horns and conceptuses were photographed. Conceptuses were dissected to isolate embryos, yolk sacs, and placentae. All embryos and placentae were photographed at equal magnification for comparison purposes.

Uteri containing placentation sites were dissected from pregnant female mice on E12.5 and E13.5 and frozen in dry ice-cooled heptane and stored at −80°C until used for histological analysis. Tissues were subsequently embedded in optimum cutting temperature (OCT) (Tissue-Tek) and were cryosectioned (10μm thick) for immunohistochemistry (IHC) studies using Leica CM-3050-S cryostat.

Placenta samples were carefully isolated, ensuring the decidual layer was peeled off. Individual samples were briefly digested in the presence of collagenase and were made into single-cell suspensions by passing them through a 40μm filter. These cell suspensions were further used for Flow analysis or FACS. Corresponding embryonic tissues were used to confirm genotypes.

For scRNA-seq, these samples were further processed using Debris Removal Solution and Dead Cell Removal Kit (Miltenyi Biotec). Red blood cell depletion from the placental suspensions were carried out using anti-Mouse Ter-119 antibody (BD Biosciences) was used.

### Flow analysis and sorting

For analyzing HSC population, placental single-cell suspensions were stained with APC-conjugated anti-mouse CD117 (c-kit) (BioLegend), PerCP/Cy5.5-conjugated anti-mouse CD34 (BioLegend), PE-conjugated anti-mouse Ly-6A/E (Sca-1) (BioLegend), PE/Cy7 anti-mouse CD45 (BioLegend) and Pacific Blue-conjugated anti-mouse Lineage cocktail (BioLegend) monoclonal antibodies. Unstained, isotype and single-color controls were used for optimal gating strategy. Samples were run on either an LSRII flow cytometer or an LSRFortessa (BD Biosciences), and the data were analyzed using FACSDiva software.

For sorting the GFP positive *Prl3d1^Cre^*+ TGCs, placental single-cell suspension from the control *Pl1^Cre^ mT/mG* and GATA Pl1-KO placentae were sorted on an LSRII flow cytometer. Placental cells from an *mT/mG* placenta were used for gating.

### Single Cell RNA-Sequencing and analysis

The transcriptomic profiles of mouse placental samples from two control (biological replicates) and two Gata2/Gata3 double knockout (biological replicate) specimens were obtained using the 10x Genomics Chromium Single Cell Gene Expression Solution (10xgenomics.com). The primary analysis of the scRNAseq data was performed using the 10x Genomics Cell Ranger pipeline (version 3.1.0). This pipeline performs sample de-multiplexing, barcode processing, and single cell 3’ gene counting. The quality of the sequenced data was assessed using the FastQC software (76). Sequenced reads were mapped to the mouse reference genome (mm10) using the STAR software (77). Individual samples were aggregated using the “cellranger aggr” tool in Cell Ranger to produce a single feature-barcode matrix containing all the sample data. This process normalizes read counts from each sample, by subsampling, to have the same effective sequencing depth. The Cell Ranger software was used to perform two-dimensional PCA and t-SNE projections of cells, and k-means clustering. The 10x Genomics Loupe Cell Browser software was used to find significant genes, cell types, and substructure within the single-cell data.

### Colony Formation Assay

Placenta cell suspensions were suspended in Iscove’s MDM with 2% fetal bovine serum and cultured using a MethoCult GF M3434 Optimum kit (STEMCELL Technologies). 10,000 cells from each sample were plated in 35-mm culture dishes (STEMCELL Technologies) and incubated at 37°C in a humidified, 5% CO2 environment for 14 days. Colonies were observed and counted using an inverted microscope.

### In Vitro Endothelial Network Assembly Assay on Matrigel

Endothelial network assembly was assayed by the formation of capillary-like structures by Human Uterine Microvascular Endothelial Cells (HUtMEC) on Matrigel (BD Biosciences). Matrigel was diluted 1:1 with a supplement-free M200 medium, poured in 12-well plates, and allowed to solidify at 37 °C. Sub confluent HUtMECs were harvested and preincubated for 1 hour in growth supplement-free M200 medium in microcentrifuge tubes. An equal volume of M200 medium containing FGF2/EGF was added. In addition, conditioned medium from *Gata2^f/f^*;*Gata3^f/f^;UBC^Cre^* trophoblast stem cells cultured in the presence (control) and absence of tamoxifen (GATA DKO) were added. The cells were plated on Matrigel (1.5 × 105 cells/well) and incubated at 37 °C and photographed at different time intervals.

### Quantitative RT-PCR

Total RNA from cells was extracted with the RNeasy Mini Kit (Qiagen). Samples isolated using FACS were further processed using the PicoPure RNA Isolation kit. cDNA samples were prepared and analyzed by qRT-PCR following procedures described earlier (78).

### Genotyping

Genomic DNA samples were prepared using tail tissues or embryonic tissues from the mice using the REDExtract-N-Amp Tissue PCR kit (Sigma-Aldrich). Genotyping was done using REDExtract-N-Amp PCR ReadyMix (Sigma-Aldrich) and respective primers. Respective primers are listed in the materials and methods section.

### Immunofluorescence

For immunostaining with mouse tissues, slides containing cryosections were dried, fixed with 4% PFA followed by permeabilization with 0.25% Triton X-100 and blocking with 10% fetal bovine serum and 0.1% Triton X-100 in PBS. Sections were incubated with primary antibodies overnight at 4 °C, washed in 0.1% Triton X-100 in PBS. After incubation (1:400, one hour, room temperature) with conjugated secondary antibodies, sections were washed, mounted using an anti-fade mounting medium (Thermo Fisher Scientific) containing DAPI and visualized using Nikon Eclipse 80i fluorescent microscope.

*mT/mG* positive embryos and cryosections were imaged directly under a Nikon Eclipse 80i fluorescent microscope.

### In situ hybridization

E13.5 whole conceptus cryosections were subjected to staining using RNAscope 2.5 HD Detection Kit (ACD Bio, Newark, CA). RNAscope probe for *Tpbpa* was used to detect the junctional zone of the mouse placenta, while hematoxylin was used as the counterstain.

### Statistical analyses

Independent data sets were analyzed by analysis of variance (ANOVA) using Student’s t-test and are presented as mean±s.e.

### Primer List

#### Primers used for genotyping

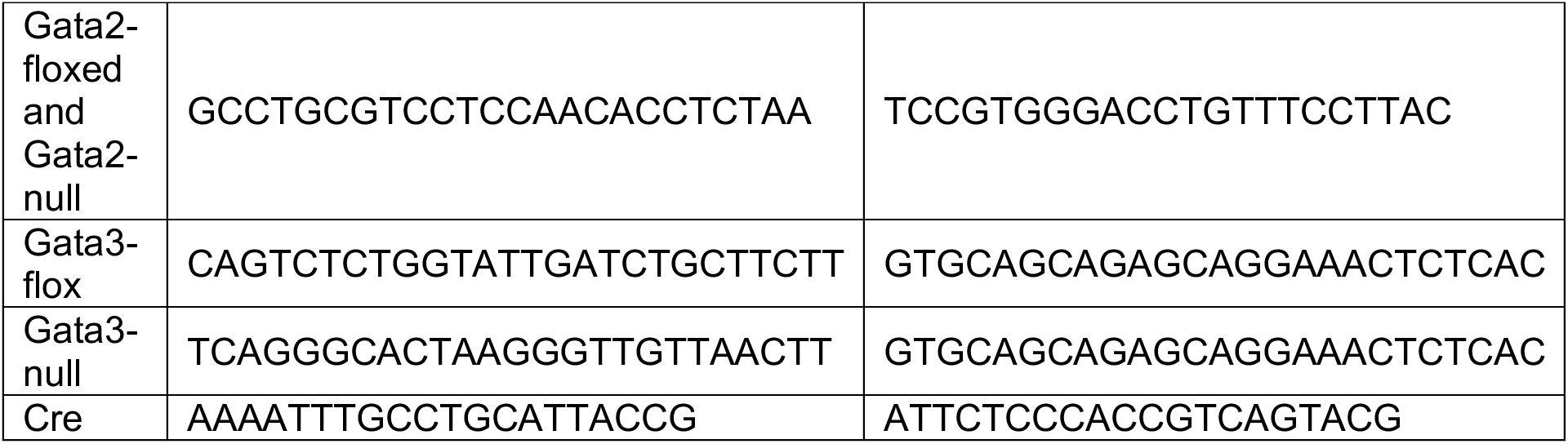

#### Primers used for quantitative RT-PCR analysis

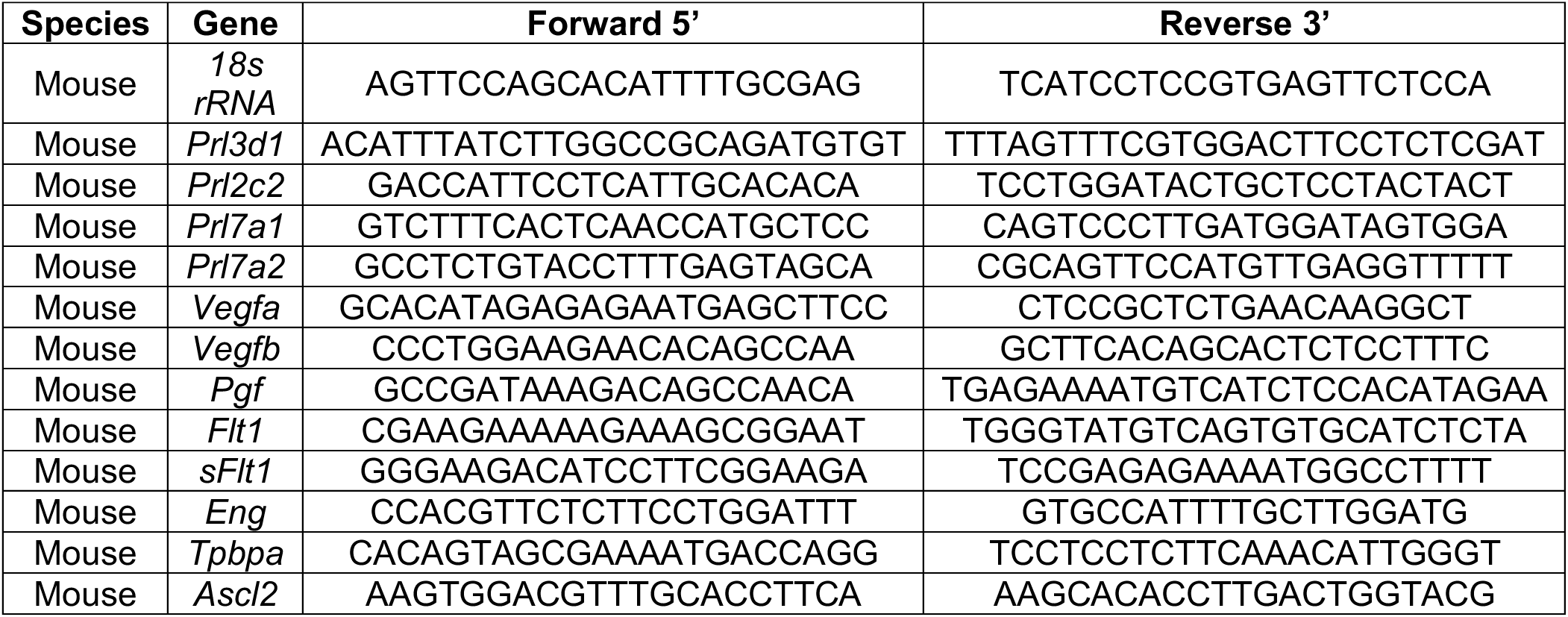

### Antibody list

#### Immunostaining

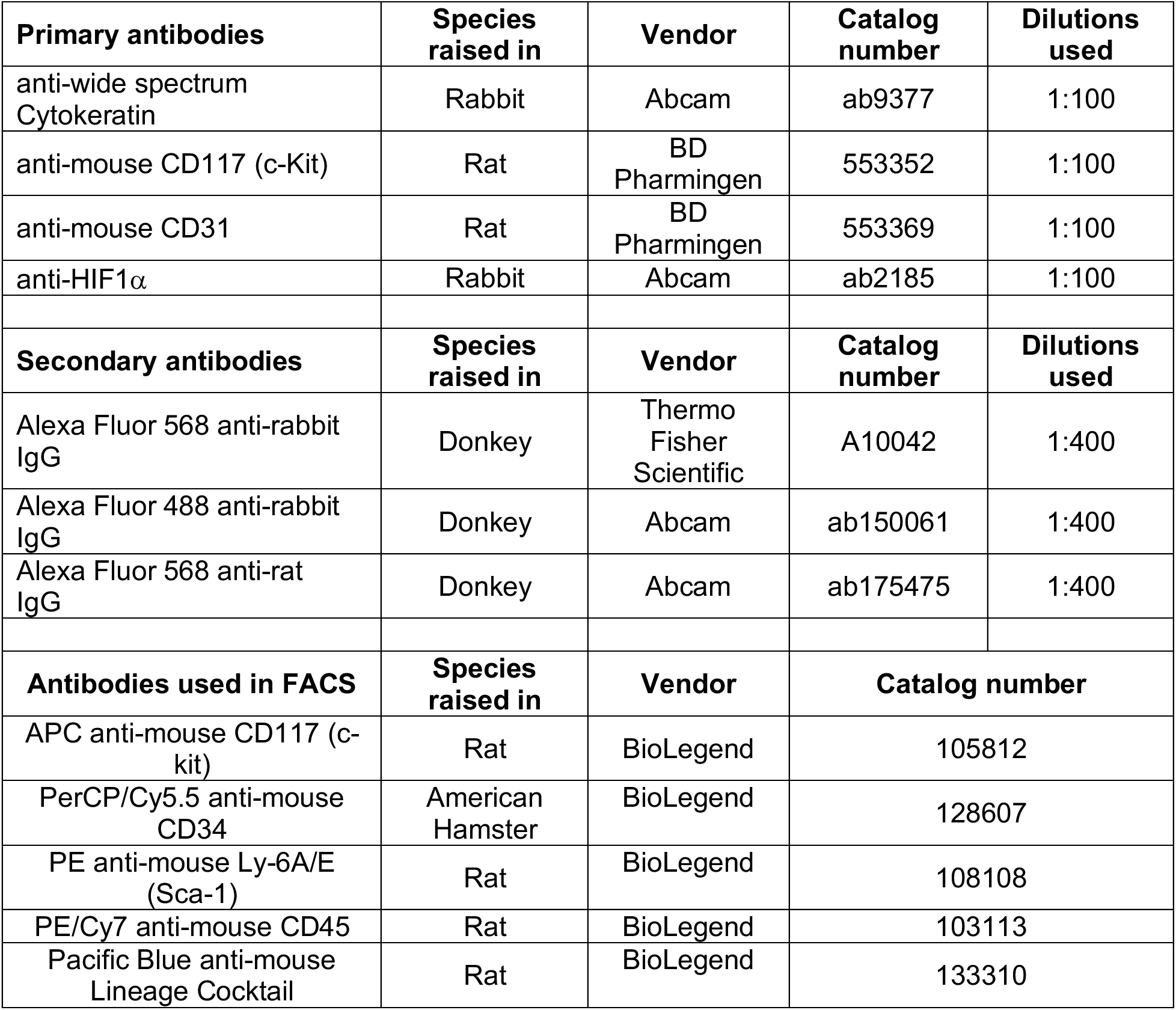

Antibody list Immunostaining
| Primary antibodies | Species raised in | Vendor | Catalog number | Dilutions used |
| --- | --- | --- | --- | --- |
| anti-wide spectrum Cytokeratin | Rabbit | Abcam | ab9377 | 1:100 |
| anti-mouse CD117 (c-Kit) | Rat | BD Pharmingen | 553352 | 1:100 |
| anti-mouse CD31 | Rat | BD Pharmingen | 553369 | 1:100 |
| anti-HIF1α | Rabbit | Abcam | ab2185 | 1:100 |
| Secondary antibodies | Species raised in | Vendor | Catalog number | Dilutions used |
| Alexa Fluor 568 anti-rabbit IgG | Donkey | Thermo Fisher Scientific | A10042 | 1:400 |
| Alexa Fluor 488 anti-rabbit IgG | Donkey | Abcam | ab150061 | 1:400 |
| Alexa Fluor 568 anti-rat IgG | Donkey | Abcam | ab175475 | 1:400 |
| Antibodies used in FACS | Species raised in | Vendor | Catalog number |  |
| APC anti-mouse CD117 (c-kit) | Rat | BioLegend | 105812 |  |
| PerCP/Cy5.5 anti-mouse CD34 | American Hamster | BioLegend | 128607 |  |
| PE anti-mouse Ly-6A/E (Sca-1) | Rat | BioLegend | 108108 |  |
| PE/Cy7 anti-mouse CD45 | Rat | BioLegend | 103113 |  |
| Pacific Blue anti-mouse Lineage Cocktail | Rat | BioLegend | 133310 |  |

## Acknowledgement

This research was supported by NIH grants HD062546, HD0098880, HD079363, a bridging grant support under the Kansas Idea Network Of Biomedical Research Excellence (K-IBRE, P20GM103418) to Soumen Paul. This study is supported by various core facilities, including the Genomics Core, the Imaging and Histology Core facility and the Bioinformatics Core of the University of Kansas Medical Center.

## Contributions

Pratik Home and Soumen Paul conceived and designed the experiments and wrote the manuscript. Pratik Home, Ananya Ghosh, Ram Parikshan Kumar, Avishek Ganguly, Bhaswati Bhattacharya, Md. Rashedul Islam and Soma Ray performed experiments. Sumedha Gunewardena performed bioinformatic analyses.

---

**Fig. S1.**
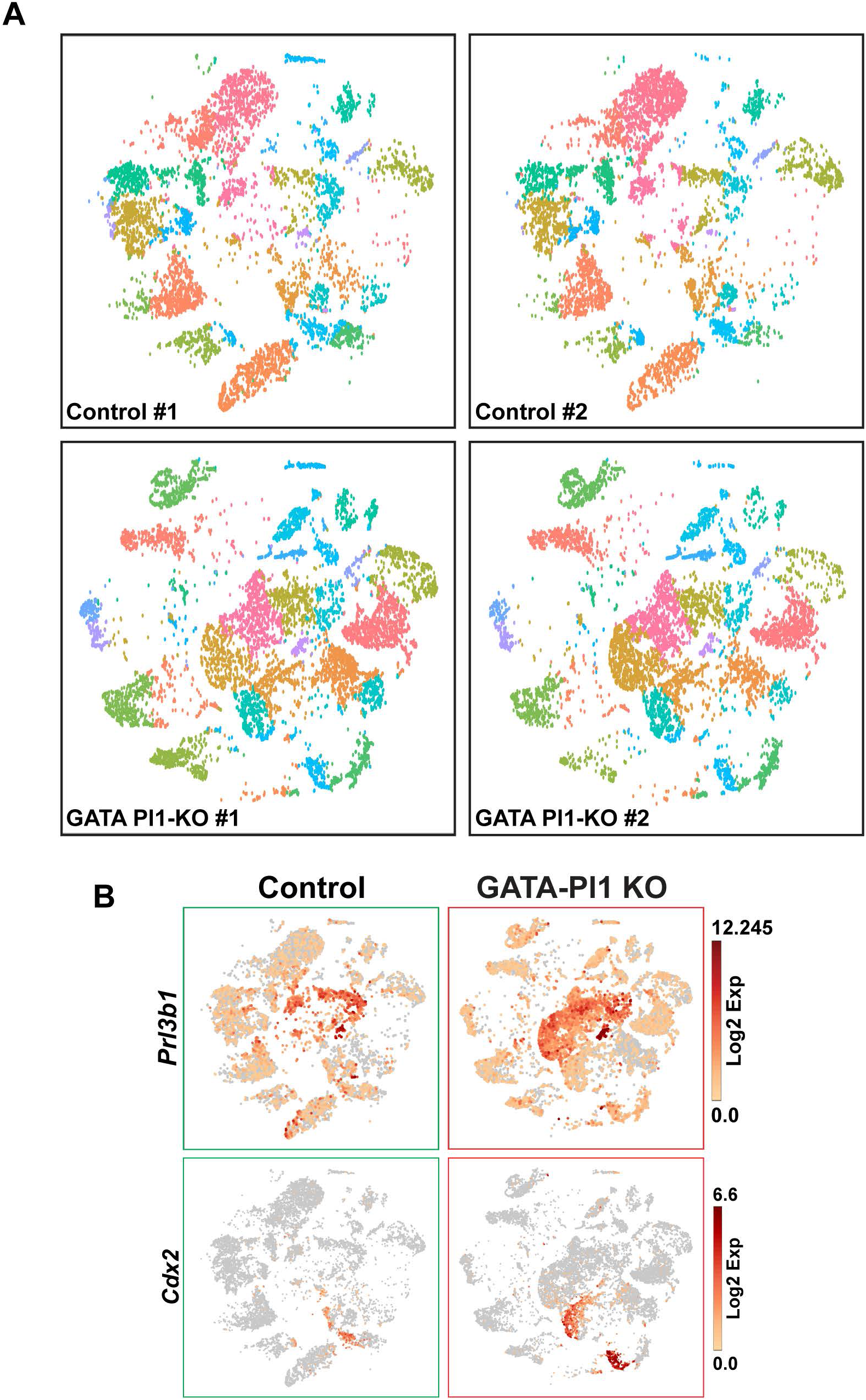
**(A)** t-SNE plot showing similar clustering patterns between two different control placental samples, and two different GATA Pl1-KO placental samples in the scRNA-seq analyses. **(B)** t-SNE plots for cells positive for *Prl3b1,* marker for both sinusoidal TGCs and parietal TGCs, and cells positive for *Cdx2* show increase in the GATA Pl-1 KO samples compared to the control.

**Fig. S2.**
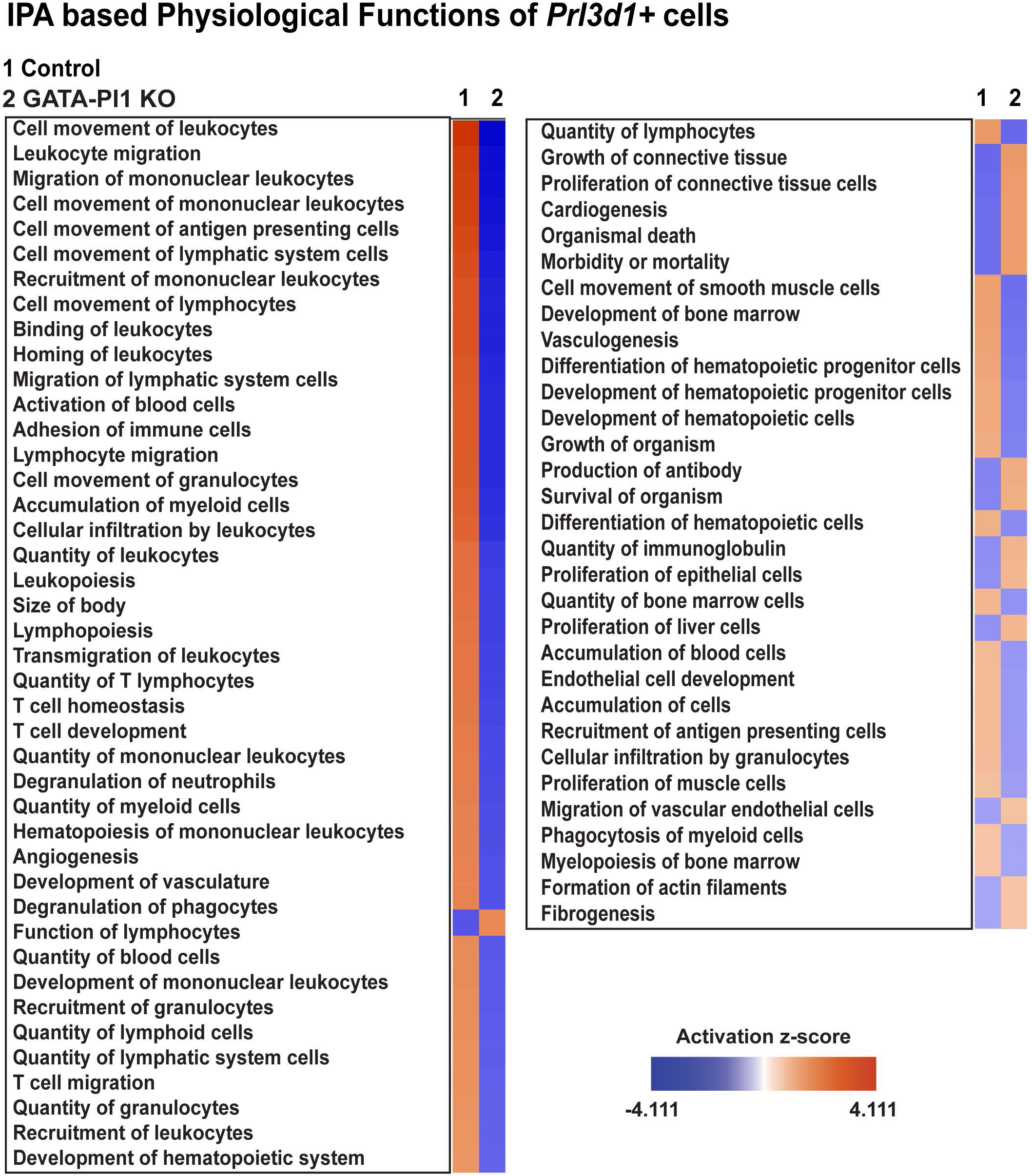
Comparison of hematopoiesis and angiogenesis related physiological functions between the *Prl3d1* positive TGCs in control and GATA-Pl1 KO samples. Significantly upregulated gene expression (p<=0.05) of the *Prl3d1* positive cells from the control and the GATA-Pl1 KO samples (compared to the rest of the cells of the corresposnding placentate) were subjected to core analysis of the Ingenuity Pathway Analysis. Physiological functions related to hematopoiesis and angiogenesis and were then compared to generate the heatmap.

**Fig. S3.**
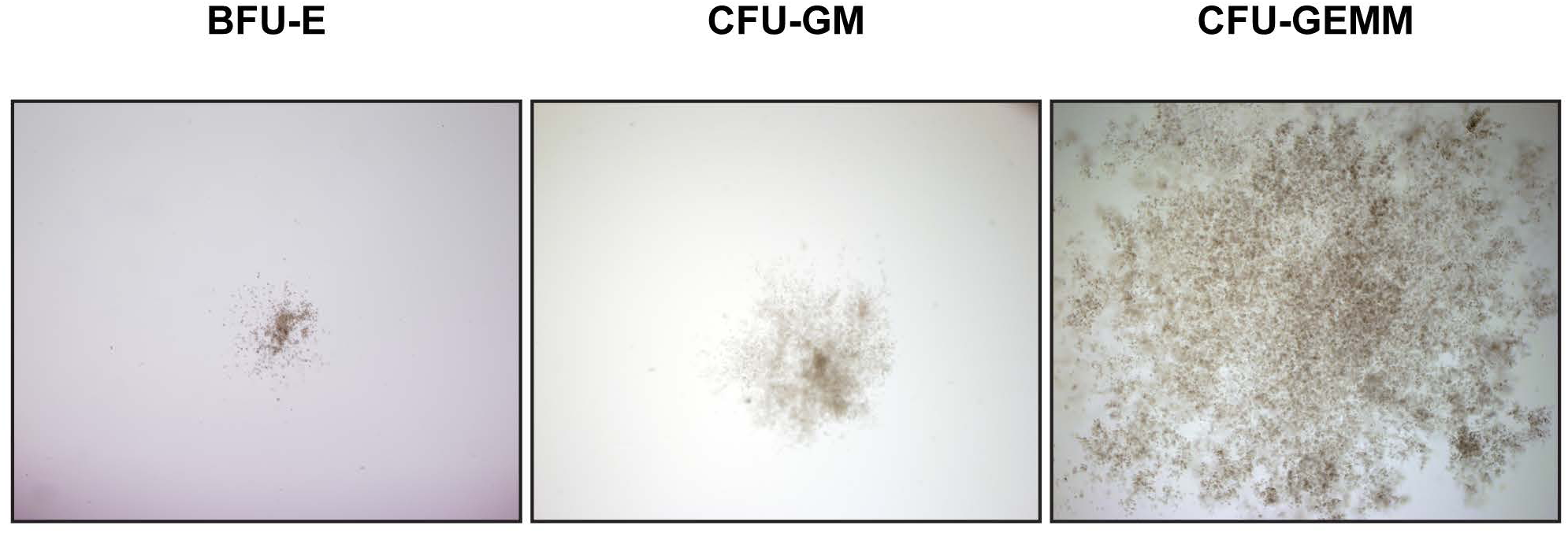
Micrographs show hematopoietic cell colonies on the methylcellulose plate. Colonies observed were mostly of the CFU-GM and CFU-GEMM nature while extremely few BFU-E colonies were present.

## References

1. L. Duley, The global impact of pre-eclampsia and eclampsia. Seminars in perinatology 33, 130–137 (2009).

2. Anonymous, Geographic variation in the incidence of hypertension in pregnancy. World Health Organization International Collaborative Study of Hypertensive Disorders of Pregnancy. American journal of obstetrics and gynecology 158, 80–83 (1988).

3. M. D. Lindheimer, J. M. Roberts, F. G. Cunningham, L. C. Chesley, Chesley’s hypertensive disorders in pregnancy (Academic Press/Elsevier, Amsterdam; Boston, ed. 3rd, 2009), pp. x, 422 p., 428 p. of plates.

4. Z. Iliodromiti et al., Endocrine, paracrine, and autocrine placental mediators in labor. Hormones (Athens) 11, 397–409 (2012).

5. S. I. Nishikawa, A complex linkage in the developmental pathway of endothelial and hematopoietic cells. Curr Opin Cell Biol 13, 673–678 (2001).

6. S. V. Koushik et al., Targeted inactivation of the sodium-calcium exchanger (Ncx1) results in the lack of a heartbeat and abnormal myofibrillar organization. FASEB J 15, 1209–1211 (2001).

7. K. E. Rhodes et al., The emergence of hematopoietic stem cells is initiated in the placental vasculature in the absence of circulation. Cell Stem Cell 2, 252–263 (2008).

8. M. Hirashima, Y. Lu, L. Byers, J. Rossant, Trophoblast expression of fms-like tyrosine kinase 1 is not required for the establishment of the maternal-fetal interface in the mouse placenta. Proceedings of the National Academy of Sciences of the United States of America 100, 15637–15642 (2003).

9. M. G. Achen, J. M. Gad, S. A. Stacker, A. F. Wilks, Placenta growth factor and vascular endothelial growth factor are co-expressed during early embryonic development. Growth Factors 15, 69–80 (1997).

10. F. Y. Tsai et al., An early haematopoietic defect in mice lacking the transcription factor GATA-2. Nature 371, 221–226 (1994).

11. P. P. Pandolfi et al., Targeted disruption of the GATA3 gene causes severe abnormalities in the nervous system and in fetal liver haematopoiesis. Nature genetics 11, 40–44 (1995).

12. S. Ray et al., Context-dependent function of regulatory elements and a switch in chromatin occupancy between GATA3 and GATA2 regulate Gata2 transcription during trophoblast differentiation. The Journal of biological chemistry 284, 4978–4988 (2009).

13. P. Home et al., GATA3 is selectively expressed in the trophectoderm of peri-implantation embryo and directly regulates Cdx2 gene expression. J Biol Chem 284, 28729–28737 (2009).

14. Y. K. Ng, K. M. George, J. D. Engel, D. I. H. Linzer, Gata Factor Activity Is Required for the Trophoblast-Specific Transcriptional Regulation of the Mouse Placental-Lactogen-I Gene. Development 120, 3257–3266 (1994).

15. G. T. Ma et al., GATA-2 and GATA-3 regulate trophoblast-specific gene expression in vivo. Development (Cambridge, England) 124, 907–914 (1997).

16. P. Home et al., Genetic redundancy of GATA factors in the extraembryonic trophoblast lineage ensures the progression of preimplantation and postimplantation mammalian development. Development 144, 876–888 (2017).

17. D. G. Simmons, J. C. Cross, Determinants of trophoblast lineage and cell subtype specification in the mouse placenta. Developmental biology 284, 12–24 (2005).

18. D. G. Simmons, A. L. Fortier, J. C. Cross, Diverse subtypes and developmental origins of trophoblast giant cells in the mouse placenta. Developmental biology 304, 567–578 (2007).

19. B. Patamalai et al., Altered expression of transforming growth factor-beta 1 mRNA and protein in mouse skin carcinogenesis. Mol Carcinog 9, 220–229 (1994).

20. M. M. Ouseph et al., Atypical E2F repressors and activators coordinate placental development. Developmental cell 22, 849–862 (2012).

21. M. D. Muzumdar, B. Tasic, K. Miyamichi, L. Li, L. Luo, A global double-fluorescent Cre reporter mouse. Genesis 45, 593–605 (2007).

22. C. Gekas et al., Hematopoietic stem cell development in the placenta. Int J Dev Biol 54, 1089–1098 (2010).

23. H. K. Mikkola, S. H. Orkin, The journey of developing hematopoietic stem cells. Development 133, 3733–3744 (2006).

24. A. H. K. El-Hashash, D. Warburton, S. J. Kimber, Genes and signals regulating murine trophoblast cell development. Mechanisms of development 127, 1–20 (2009).

25. B. M. Bany, J. C. Cross, Post-implantation mouse conceptuses produce paracrine signals that regulate the uterine endometrium undergoing decidualization. Developmental biology 294, 445–456 (2006).

26. Y. Hu et al., IFN-gamma-mediated extravillous trophoblast outgrowth inhibition in first trimester explant culture: a role for insulin-like growth factors. Mol Hum Reprod 14, 281–289 (2008).

27. E. W. Carney, V. Prideaux, S. J. Lye, J. Rossant, Progressive expression of trophoblast-specific genes during formation of mouse trophoblast giant cells in vitro. Molecular reproduction and development 34, 357–368 (1993).

28. T. Parisi et al., Cyclins E1 and E2 are required for endoreplication in placental trophoblast giant cells. The EMBO journal 22, 4794–4803 (2003).

29. L. Anson-Cartwright et al., The glial cells missing-1 protein is essential for branching morphogenesis in the chorioallantoic placenta. Nature genetics 25, 311–314 (2000).

30. E. Basyuk et al., Murine Gcm1 gene is expressed in a subset of placental trophoblast cells. Developmental dynamics : an official publication of the American Association of Anatomists 214, 303–311 (1999).

31. M. Muntener, Y. C. Hsu, Development of trophoblast and placenta of the mouse. A reinvestigation with regard to the in vitro culture of mouse trophoblast and placenta. Acta Anat (Basel) 98, 241–252 (1977).

32. D. I. Linzer, S. J. Fisher, The placenta and the prolactin family of hormones: regulation of the physiology of pregnancy. Mol Endocrinol 13, 837–840 (1999).

33. S. Yotsumoto et al., Expression of adrenomedullin, a hypotensive peptide, in the trophoblast giant cells at the embryo implantation site in mouse. Developmental biology 203, 264–275 (1998).

34. H. WeilerGuettler, W. C. Aird, H. Rayburn, M. Husain, R. D. Rosenberg, Developmentally regulated gene expression of thrombomodulin in postimplantation mouse embryos. Development 122, 2271–2281 (1996).

35. S. J. Lee, F. Talamantes, E. Wilder, D. I. H. Linzer, D. Nathans, Trophoblastic Giant-Cells of the Mouse Placenta as the Site of Proliferin Synthesis. Endocrinology 122, 1761–1768 (1988).

36. J. C. Cross et al., Trophoblast functions, angiogenesis and remodeling of the maternal vasculature in the placenta. Mol Cell Endocrinol 187, 207–212 (2002).

37. K. Bollerot, C. Pouget, T. Jaffredo, The embryonic origins of hematopoietic stem cells: a tale of hemangioblast and hemogenic endothelium. APMIS 113, 790–803 (2005).

38. C. Gekas, F. Dieterlen-Lievre, S. H. Orkin, H. K. A. Mikkola, The placenta is a niche for hematopoietic stem cells. Developmental cell 8, 365–375 (2005).

39. K. Ottersbach, E. Dzierzak, The murine placenta contains hematopoietic stem cells within the vascular labyrinth region. Developmental cell 8, 377–387 (2005).

40. R. A. Smith, C. A. Glomski, “Hemogenic endothelium” of the embryonic aorta: Does it exist? Dev Comp Immunol 6, 359–368 (1982).

41. J. Palis, M. C. Yoder, Yolk-sac hematopoiesis: the first blood cells of mouse and man. Exp Hematol 29, 927–936 (2001).

42. F. Shalaby et al., Failure of blood-island formation and vasculogenesis in Flk-1-deficient mice. Nature 376, 62–66 (1995).

43. F. Shalaby et al., A requirement for Flk1 in primitive and definitive hematopoiesis and vasculogenesis. Cell 89, 981–990 (1997).

44. K. Choi, M. Kennedy, A. Kazarov, J. C. Papadimitriou, G. Keller, A common precursor for hematopoietic and endothelial cells. Development 125, 725–732 (1998).

45. I. Kim, O. H. Yilmaz, S. J. Morrison, CD144 (VE-cadherin) is transiently expressed by fetal liver hematopoietic stem cells. Blood 106, 903–905 (2005).

46. R. Nasrallah et al., Identification of novel regulators of developmental hematopoiesis using Endoglin regulatory elements as molecular probes. Blood 128, 1928–1939 (2016).

47. A. Kubo et al., The homeobox gene HEX regulates proliferation and differentiation of hemangioblasts and endothelial cells during ES cell differentiation. Blood 105, 4590–4597 (2005).

48. M. A. Park et al., Activation of the Arterial Program Drives Development of Definitive Hemogenic Endothelium with Lymphoid Potential. Cell Rep 23, 2467–2481 (2018).

49. K. L. Marcelo, L. C. Goldie, K. K. Hirschi, Regulation of endothelial cell differentiation and specification. Circ Res 112, 1272–1287 (2013).

50. T. E. North et al., Runx1 expression marks long-term repopulating hematopoietic stem cells in the midgestation mouse embryo. Immunity 16, 661–672 (2002).

51. W. Risau, Embryonic angiogenesis factors. Pharmacol Ther 51, 371–376 (1991).

52. J. Wilting, B. Christ, Embryonic angiogenesis: a review. Naturwissenschaften 83, 153–164 (1996).

53. G. Breier, Angiogenesis in embryonic development--a review. Placenta 21 Suppl A, S11–15 (2000).

54. C. J. Drake, P. A. Fleming, Vasculogenesis in the day 6.5 to 9.5 mouse embryo. Blood 95, 1671–1679 (2000).

55. P. Carmeliet et al., Synergism between vascular endothelial growth factor and placental growth factor contributes to angiogenesis and plasma extravasation in pathological conditions. Nature medicine 7, 575–583 (2001).

56. M. Shibuya, Vascular Endothelial Growth Factor (VEGF) and Its Receptor (VEGFR) Signaling in Angiogenesis: A Crucial Target for Anti- and Pro-Angiogenic Therapies. Genes Cancer 2, 1097–1105 (2011).

57. J. C. Chappell et al., Flt-1 (VEGFR-1) coordinates discrete stages of blood vessel formation. Cardiovasc Res 111, 84–93 (2016).

58. S. Banerjee, S. K. Dhara, M. Bacanamwo, Endoglin is a novel endothelial cell specification gene. Stem Cell Res 8, 85–96 (2012).

59. K. D. C. Dahl et al., Hypoxia-indulcible factors 1 alpha and 2 alpha regulate trophoblast differentiation. Molecular and cellular biology 25, 10479–10491 (2005).

60. D. M. Adelman, M. Gertsenstein, A. Nagy, M. C. Simon, E. Maltepe, Placental cell fates are regulated in vivo by HIF-mediated hypoxia responses. Gene Dev 14, 3191–3203 (2000).

61. D. M. Adelman, E. Maltepe, M. C. Simon, Multilineage embryonic hematopoiesis requires hypoxic ARNT activity. Blood 94, 254a–254a (1999).

62. E. Maltepe, J. V. Schmidt, D. Baunoch, C. A. Bradfield, M. C. Simon, Abnormal angiogenesis and responses to glucose and oxygen deprivation in mice lacking the protein ARNT. Nature 386, 403–407 (1997).

63. R. Tal et al., Effects of hypoxia-inducible factor-1alpha overexpression in pregnant mice: possible implications for preeclampsia and intrauterine growth restriction. The American journal of pathology 177, 2950–2962 (2010).

64. K. D. Cowden Dahl et al., Hypoxia-inducible factors 1alpha and 2alpha regulate trophoblast differentiation. Mol Cell Biol 25, 10479–10491 (2005).

65. D. M. Adelman, M. Gertsenstein, A. Nagy, M. C. Simon, E. Maltepe, Placental cell fates are regulated in vivo by HIF-mediated hypoxia responses. Genes Dev 14, 3191–3203 (2000).

66. D. Hu, J. C. Cross, Development and function of trophoblast giant cells in the rodent placenta. Int J Dev Biol 54, 341–354 (2010).

67. G. T. Ma et al., GATA-2 and GATA-3 regulate trophoblast-specific gene expression in vivo. Development 124, 907–914 (1997).

68. B. Zhou, X. Kong, D. I. Linzer, Enhanced recovery from thrombocytopenia and neutropenia in mice constitutively expressing a placental hematopoietic cytokine. Endocrinology 146, 64–70 (2005).

69. S. Bhattacharyya, J. Lin, D. I. Linzer, Reactivation of a hematopoietic endocrine program of pregnancy contributes to recovery from thrombocytopenia. Mol Endocrinol 16, 1386–1393 (2002).

70. S. Yotsumoto et al., Expression of adrenomedullin, a hypotensive peptide, in the trophoblast giant cells at the embryo implantation site in mouse. Dev Biol 203, 264–275 (1998).

71. M. Li et al., Fetal-derived adrenomedullin mediates the innate immune milieu of the placenta. J Clin Invest 123, 2408–2420 (2013).

72. K. Kanenishi, H. Kuwabara, M. Ueno, H. Sakamoto, T. Hata, Immunohistochemical adrenomedullin expression is decreased in the placenta from pregnancies with pre-eclampsia. Pathol Int 50, 536–540 (2000).

73. S. Tanaka, T. Kunath, A. K. Hadjantonakis, A. Nagy, J. Rossant, Promotion of trophoblast stem cell proliferation by FGF4. Science 282, 2072–2075 (1998).

74. M. A. Charles et al., Pituitary-specific Gata2 knockout: effects on gonadotrope and thyrotrope function. Mol Endocrinol 20, 1366–1377 (2006).

75. J. Zhu et al., Conditional deletion of Gata3 shows its essential function in T(H)1-T(H)2 responses. Nature Immunology 5, 1157–1165 (2004).

76. S. Andrews, FastQC: a quality control tool for high throughput sequence data. Available online at: http://www.bioinformatics.babraham.ac.uk/projects/fastqc. (2010).

77. A. Dobin et al., STAR: ultrafast universal RNA-seq aligner. Bioinformatics 29, 15–21 (2013).

78. D. Dutta, S. Ray, J. L. Vivian, S. Paul, Activation of the VEGFR1 chromatin domain: an angiogenic signal-ETS1/HIF-2alpha regulatory axis. The Journal of biological chemistry 283, 25404–25413 (2008).

